# Neurexin and Neuroligin-based adhesion complexes drive axonal arborisation growth independent of synaptic activity

**DOI:** 10.1101/182808

**Authors:** William D Constance, Amrita Mukherjee, Yvette E Fisher, Sînziana Pop, Eric Blanc, Yusuke Toyama, Darren W Williams

## Abstract

Building arborisations of the right size and shape is fundamental for neural network function. Live imaging studies in vertebrate brains strongly suggest that nascent synapses are critical for branch growth during the development of axonal and dendritic arborisations. The molecular mechanisms underlying such ‘synaptotropic’ events are largely unknown.

Here we present a novel system in *Drosophila* for studying the development of complex axonal arborisations live, *in vivo* during metamorphosis. In these growing axonal arborisations we see a relationship between the punctate localisations of presynaptic components and branch dynamics that is very similar to synaptotropic growth described in fish and frogs. These presynaptic components however do not appear to represent functional presynaptic release sites and are not paired with clusters of neurotransmitter receptors. Pharmacological and genetic knockdowns of evoked and spontaneous neurotransmission do not impact the outgrowth of these neurons. Instead, we find that axonal branch growth is regulated by the dynamic focal localisations of synaptic adhesion proteins Neurexin and Neuroligin. These adhesion complexes provide selective stability for filopodia by a ‘stick and grow’-based mechanism wholly independent of synaptic activity.

## Introduction

Neurons are the most structurally diverse and complex cell type we know of (Bullock and Horridge, 1965). Their tree-like arborisations are critical for collecting, integrating and disseminating information between different synaptic partners. During development, arborisations grow dynamically and this morphogenesis sets limits on their final location, possible synaptic partners and core electrophysiological properties (Chagnac-Amatai *et al.*, 1990; Roberts *et al.*, 2014). Although there are recognisable cell-type specific shapes, each individual neuron has its own unique pattern of branching and connectivity. The ‘generative rules’ for constructing complex arborisations are believed to be encoded by genetic algorithms that play out in different developmental contexts to produce distinct morphological types (Cuntz at al., 2010; Teeter & Stevens., 2011; Chen & Haas, 2011; Hassan & Heisinger, 2015). Which molecules and mechanisms underlie this remains a major unanswered question within neuroscience.

From observations on the earliest phases of motoneuron dendrite growth in the spinal cord Vaughn and colleagues forwarded the *synaptotropic hypothesis* (Vaughn *et al.*, 1974 &. The *synaptotropic hypothesis* posits that the stability of growing axonal and dendritic branches is controlled by the selective stabilisation of processes by nascent synapses, encouraging growth into territories rich in potential synaptic partners. Berry and colleagues, working on Purkinje cells, forwarded the same idea, calling it the *synaptogenic filopodial theory* (Berry & Bradley, 1976). Since then, live imaging in vertebrate brains has revealed dynamic axonal and dendritic growth (Kaethner & Stuermer, 1992; Wu *et al.*, 1999; Jontes *et al.*, 2000; Hossain *et al.*, 2012) where the arrival and localisation of synaptic machineries is correlated with branch stabilisation (Alsina *et al.*, 2001; Niell *et al.*, 2004; Meyer & Smith, 2006; Ruthazer *et al.*, 2006). Data on *Xenopus* tectal neuron dendritic growth revealed a role for Neurexin (Nrx) and Neuroligin (Nlg) in branch dynamics, where Nrx-Nlg interactions are believed to direct the genesis and maturation of opposing hemisynapses, after which neurotransmission stabilises branches (Chen *et al.*, 2010). The idea that synaptic transmission stabilises nascent contacts during elaboration is supported by a number of studies (Rajan *et al* 1999; Sin *et al.*, 2002; Ruthazer *et al.*, 2003; Haas *et al.*, 2006; Ruthazer *et al.*, 2006). In contrast, other work has suggested that activity has little impact on large-scale features of tree growth but plays a more nuanced role in refining structural connectivity during activity dependent plasticity (Verhage *et al.*, 2000; Varoquoaeux *et al.*, 2002; Hua *et al.*, 2005; Fredj *et al.*, 2010).

Two key questions arise from this work: firstly, are growing branches stabilised by local ‘synaptogenic’ events? Secondly, are such nascent synapses being ‘use-tested’ by conventional neurotransmission? Here we describe a new system that we have pioneered in *Drosophila* to explore these questions. Unlike in the fly embryo where arborisations are very small and built very rapidly (<4 hours), our system takes advantage of the large axonal arborisations of the pleural muscle motoneurons (PM-Mns) that are built over an extended period, during metamorphosis. Also, unlike the growth of larval motoneuron terminals, which grow incrementally to scale with the changes in muscle size during larval life, the PM-Mn axonal arborisations grow exuberantly, by trial-and-error, very much like complex neurons found in vertebrate central nervous systems.

During development we see a consistent relationship between the distribution of presynaptic machineries and branch dynamics similar to that found in the retinal ganglion cell axons in fish and frogs. Importantly, we find that branch growth is driven by dynamic complexes in synaptic partners that we term neuritic adhesion complexes (NACs). These NACs contain Nlg1 (postsynaptically), along with Nrx, Syd1 and Liprin-α (presynaptically), and act locally to stabilise filopodia by a ‘*stick and grow*’ mechanism without the need of synaptic vesicle release machinery or functional synapses.

## Results

### Establishing a new model for exploring complex arborisation growth

In a search for a system to study complex arbor growth live, *in vivo* we identified the axonal arborisations of the motoneurons that innervate the abdominal pleural muscles of the adult fly. We refer to these as the pleural muscle motoneurons (PM-Mns), the axonal arborisations of which are large, complex and easily accessible throughout metamorphosis. Importantly, the PM-Mn system allows one to genetically manipulate either synaptic partner whilst independently imaging the other.

Each adult abdominal hemisegment (from A1-A7) contains a pair of motoneurons that exit the nerve onto the lateral body wall. In the adult, each motoneuron forms an arborisation that innervates between 15 and 18 parallel muscle fibres that span dorso-ventrally, from the tergites to sternites (Figures 1A&B). In contrast to the rigid, target-specificity found in the larval neuromuscular system, the innervation of the pleural muscles shows variation between segments and between animals. Although no one-to-one neuron/muscle targeting is found, arborisations achieve a consistent spatial organisation with regularly sized, non-overlapping projection fields.

**Figure 1.**
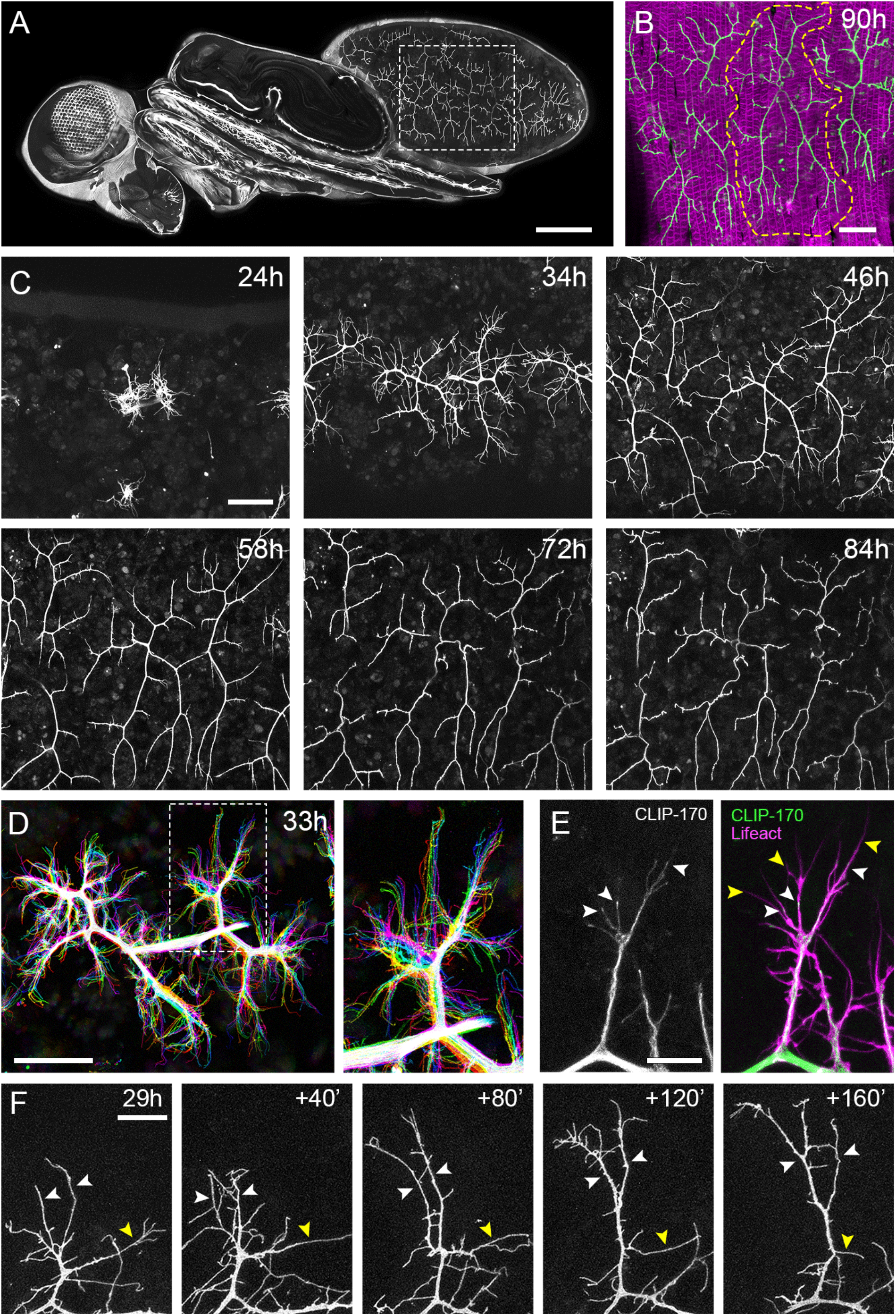
The pleural motoneurons show dynamic growth during development. (**A**) Tiled confocal stacks show a dorso-lateral view of a pupa at 90h APF expressing *UAS-myr::GFP* in the pattern of the glutamatergic driver *OK371-GAL4*. Dashed lines demarcate arborisation innervating segments A3&4 (**B**) Muscles labelled with *mCD8::Cherry (Mef2-GAL4*) (magenta) and glutamatergic neurons labelled with *myr::GFP (VGlut-LexA* (Diao *et al*., 2015)) (green) show the innervation of the pleural muscles in segments A3&4 at 90h APF. Dashed lines demarcate the anterior arborisation in 4A (A4A). (**C**) A time series follows the growth of the same arborisations in segment A3 from 24h to 84h APF in 10-14h intervals (*VGlut-LexA>myr::GFP)*. (**D**) A temporally colour coded projection of a 10 minute time-lapse with 2 minute intervals demonstrates the dynamics of a pair of arborisations in segment A3 at 32h APF. (**E**) Subcellular localisations of cytoskeletal components offer a molecular basis for delineating arbor features. Microtubules, revealed by CLIP-170, are concentrated in branches and dynamically invade nascent branchlets (white arrowheads). Filopodia (yellow arrowheads), are rich with actin, revealed by Lifeact::Ruby, but largely devoid of microtubules. (**F**) A time series with 40-minute intervals shows the growth of a branch at 29h APF. White arrowheads indicate filopodia which are stabilised and mature into branches. The yellow arrowhead indicates a branch which regresses into a filopodium. Scale bars: 250*μ*m (**A**), 50*μ*m (**B,C,D**), 10*μ*m (**E**), 20*μ*m (**F**).

To gain insight into the global development of the PM-Mn axon terminals, we imaged hemi-segment A3 from 24h to 84h after puparium formation (APF) (Figure 1C). By 24h APF the majority of larval muscles and neurons that innervate them are removed by programmed cell death and phagocytosis (Currie & Bate, 1991). The dorsally projecting peripheral nerve maintains four contacts with the epidermis during pupariation and allows the continued innervation of the persistent larval muscles. At around 24h APF axonal regrowth commences with the emergence of filopodia-rich growth cones from the nerve. By 34h APF primary branches are established and festooned with filopodia. The two PM-Mns within each hemisegment segregate into anterior and posterior domains and increase in size and complexity until approximately 72h APF. Following this, there is a period of maturation during which varicosities form along all but the most proximal branches. By 84h APF the morphology of the arborisations is indistinguishable from that observed at eclosion.

Live imaging at 2-minute intervals revealed that branch growth is highly dynamic, involving large numbers of filopodia which continually explore the local environment (Figure 1D). This exploration takes place almost exclusively within the plane of the developing muscles. Throughout this article, we define and refer to filopodia and branches as follows. According to anatomical and molecular criteria, filopodia are considered to be thin (<0.3*μ*m), motile protrusions with cytoskeletons comprised of parallel F-actin filaments (Mattila & Lappalainen, 2008). In contrast, branches are more voluminous, of higher calibre and less dynamic, with cores containing parallel microtubules and noticeably less F-actin. Using Lifeact, a fluorescently tagged cytoskeletal probe that reveals F-actin (Riedl *et al.*, 2008), we see an enrichment in thin ‘filopodia-like’ structures (Figure 1E). In contrast, the plus-end microtubule binding protein CLIP-170 (Stramer *et al.*, 2010), that reveals microtubules, is found in what we refer to as branches but is mostly excluded from filopodia. Growing neurites are dynamic and constantly changing, but every effort has been made to be consistent with our classification.

Longer imaging intervals revealed the more substantial structural changes to arborisations during early phases of growth (Figure 1F). This showed us that the majority of filopodia are transient since large numbers are generated and lost between frames. Despite this, a small number of filopodia persist, pioneering individual branches (white arrowheads). In addition to construction, branches also collapse back into single filopodia (yellow arrowhead), highlighting the instability and continuous remodelling of the arborisations at this early stage (see movie 1).

### The distribution of presynaptic components during axonal branch growth

To explore the growth of PM-Mn axonal arborisations and their muscle targets we imaged both the neuron and muscle progenitors (myoblasts) during the earliest phases of outgrowth. At 35h APF neurons and myoblasts are directly apposed to one another (Figure 2A and movie 2). Growing axons are enveloped by clusters of myoblasts, while the more distal growth cones and filopodia extend over sheets of immature myotubes.

**Figure 2.**
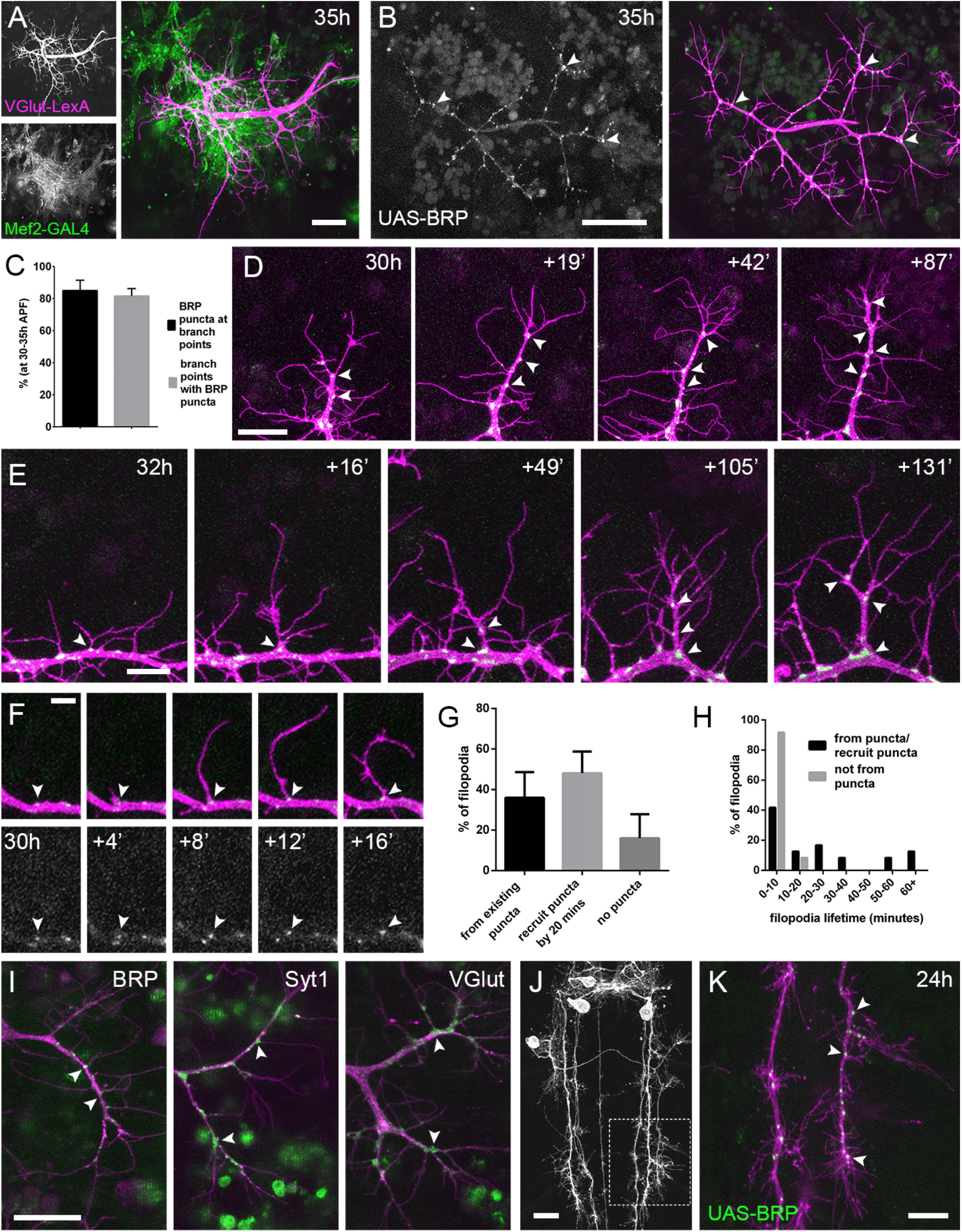
Arrival and localisation of presynaptic components is tightly linked with arbor growth. (**A**) Muscles labelled with *myr::tdTomato (Mef2-GAL4* (Ranganayakulu *et al.*, 1996)) and motoneurons labelled with *myr::GFP (VGlut-LexA*) at 35h APF show the intimate development of the pleural arborisations and their postsynaptic targets. (**B**) *BRP::RFP* expressed in motoneurons (*OK371-GAL4*) (green) decorates a pair of axon terminals (labelled also with myr::GFP) (magenta) at 35h APF, particularly at branch points (arrowheads). (**C**) At stages 30-35h APF 85.05 ± 6.42% of BRP::RFP puncta are localised at branch points or the bases of filopodia and 81.62 ± 4.64% of branch points/bases of filopodia host a punctum (n = 5). (**D**) BRP::RFP puncta form in the wake of growing axon branches. Time series shows the rapid arrival of BRP::RFP puncta, indicated by arrowheads, in the growth cone of an extending branch, producing a new segment studded with BRP::RFP puncta, most of which mark sites of filopodia growth. (**E**) Time series shows the maturation of a filopodium which originates from a BRP::RFP punctum (indicated by the arrowhead in the first frame) into a branch. Additional BRP::RFP puncta, indicated by arrowheads, are rapidly recruited to new branch nodes (**F**) Time series shows the emergence of a filopodium from a site marked by a BRP::RFP punctum (arrowheads). (**G**) The proportions of filopodia which (i) emerge from existing puncta, (ii) recruit a punctum to their base within 20 minutes and (iii) do not recruit puncta to their bases (83 filopodia from 3 time-lapse movies at stages 30-33h APF). (**H**) Lifetimes of filopodia which originate from/recruit puncta within 20 minutes of emergence (21.67 ± 20.30 mins, n = 24) or do not recruit puncta (3.83 ± 2.76 mins, n = 24) in a histogram with 20 minute bins. Filopodia which originate from puncta/recruit puncta are significantly longer lived than those that do not recruit puncta (Mann-Whitney U = 73, p < 0.0001, two-tailed). (**I**) Protein trap lines reveal the localisations of endogenous BRP (MiMIC collection (Venken *et al*., 2011)), Syt1 and VGlut (CPTI collection (Lye *et al*., 2014)) (arrowheads) in arborisations staged between 30h and 33h APF. (**J&K**) Like in motoneurons, *BRP::RFP* expressed in a central, *eve+* interneuron (*Rn2-FLPout*) has a punctate distribution in the growing output arborisations at points of branch and filopodia growth, shown by arrowheads, at 24h APF. The region marked by dashed lines in J is expanded in K. Bars represent SDs. Scale bars: 20*μ*m (**A,D,I,J**), 50*μ*m (**B**), 10*μ*m (**E,K**), 5*μ*m (**F**).

In light of this very close association between the synaptic partners, we asked whether presynaptic components are present in the distal branches at this early stage, since previous work in vertebrates has forwarded that nascent synapses play a key role in branch stabilisation. Using an RFP tagged version of Bruchpilot (BRP), a homolog of ELKS/CAST, expressed with *OK371-GAL4* (Mahr & Aberle., 2006), we see distinct, punctate distributions in axon branches, with accumulations at branch nodes and at the base of filopodia (Figure 2B). An analysis of BRP::RFP puncta in axon terminals of 5 animals staged from 30h to 35h APF revealed that 85.1 ± 6.4% (SD) of total BRP puncta localised at branch points and the base of filopodia, and 81.6 ± 4.6% of branch points/bases had a punctum (Figure 2C).

To look at the dynamics of BRP::RFP and branching we imaged PM-Mn arborisations at 2-minute intervals. This showed that BRP::RFP puncta are rapidly recruited to the tips of growing branches where they mark the sites of new filopodia growth (Figure 2D). Successive rounds of filopodia extension and stabilisation in this manner produce branches studded with BRP::RFP puncta. Filopodia which are generated from these puncta can give rise to new branches (Figure 2E).

The dynamics of BRP::RFP puncta in these time-lapse sequences speaks to a relationship between arbor growth and the distribution of presynaptic proteins (Figure 2F). To quantify this we analysed 83 filopodia generation events from 3 time-lapse movies of arborisations staged between 30h and 33h APF. Our analysis showed that 36.0 ± 12.6% of filopodia emerged from existing BRP puncta, 48.1 ± 10.7% recruited a punctum within 20 minutes of their formation and 16.0 ± 11.9% failed to recruit an obvious punctum at all (Figure 2G). The frequency of filopodia emerging from or recruiting a BRP::RFP punctum 71.4 ± 11.0% was significantly greater than a conservative expected frequency of 50% *(X^2^* = 36.4, p < 0.0001). In addition, we found that 100% of filopodia that failed to recruit puncta to their bases were eliminated within 20 minutes of their emergence (n = 24) (Figure 2H). In contrast, although 54.2% of filopodia that originate from, or recruited BRP::RFP puncta were also lost within 20 minutes, 33.3% survived between 20 and 60 minutes and a further 12.5% persisted for over an hour (n = 24). Thus, the lifetimes of filopodia associated with puncta at their bases were significantly longer than those without (Mann-two-tailed Whitney U = 73, p < 0.0001) (See movie 3).

To verify the accuracy of GAL4 driven BRP::RFP, we imaged endogenous BRP protein tagged with GFP (Figure 2I) as immunohistochemistry is particularly difficult on fragile, early pupal body walls. The BRP::GFP protein trap revealed a punctate distribution at branch points and base of filopodia, as we had seen with the BRP::RFP reporter. Other GFP tagged protein trap lines that label synaptic vesicle (SV) associated proteins Syt1 and VGlut showed less punctate localisations, yet were still clearly concentrated in axon branches at the base of filopodia (Fig 2I). Interestingly we never saw these proteins within filopodia.

We wondered whether this node localisation of pre-synaptic machinery in the growing branches was specific to PM-Mns or is more widespread in the fly nervous system. To test this we imaged the output terminals of *Eve* (*Even-skipped*) positive interneurons in the thoracic neuromeres during metamorphosis (24h APF) (Roy *et al.*, 2007). In these we found very similar distributions to those of the PM-Mns, with 79.5 ± 3.7% of puncta found at branch points or the base of filopodia (n = 8 output fields from 4 individuals) (Figures 2J&K).

### PM-Mns elaborate their axonal arborisations without synaptic activity

The localisation and dynamics of presynaptic machineries described above mirrors that seen in *Xenopus* and zebrafish tecta and suggested to us a possible role for synapses and synaptic activity in branch growth during PM-Mn elaboration (Alsina *et al.*, 2001; Niell *et al.*, 2004; Meyer & Smith, 2006; Ruthazer *et al.*, 2006).

To explore the role of neural activity in arborisation growth we mapped the electrical development of the PM-Mns using the genetically encoded calcium indicator *GCaMP6m* as a proxy (Movies 4-6). Calcium activity, recorded as changes in fluorescence, is shown as heat-registered kymographs at different stages of development (Figure 3A). The absence of any overt changes in fluorescence at 32h APF indicates that at this stage the motoneurons are electrically inactive. It was not until 42h APF that we saw the first calcium events indicative of membrane depolarisations. These events were isolated and infrequent. The frequency of calcium events increased sharply and by 48h APF activity was characterised by episodes of transients in quick succession lasting around 8 minutes, interspersed with longer periods of quiescence. Approaching the final stages of arbor growth, at 74h APF activity continued to be defined by similar periods of activity and quiescence, but at these stages bouts contained far greater numbers of events.

**Figure 3.**
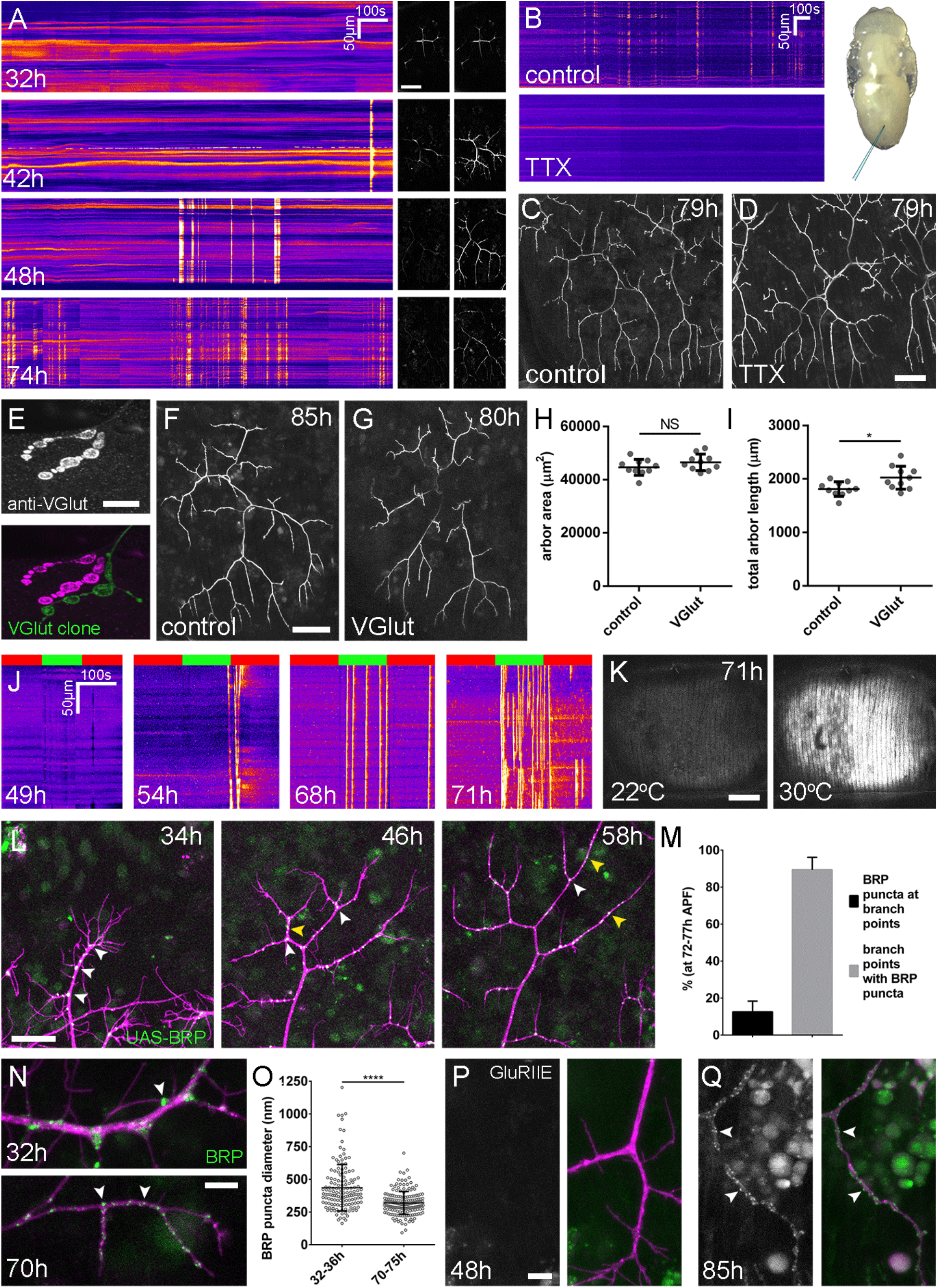
Arbor growth takes place without the formation of functional synapses. (**A**) Heat registered kymographs show calcium activity in motoneuron axon terminals at 32, 42, 48 and 74h APF, measured by changes in GCaMP6m fluorescence (Δf) (*OK371-GAL4).* Sequential images show examples of Δfs. (**B**) Kymographs display GCaMP6m Δfs in motoneuron axons at 78h APF in a pupa injected with a control solution of PBS (top) and a solution of 0.5mM TTX in PBS (bottom) into the abdomen as indicated in the image of a pupa. (**C**) Arborisations in segment A3 (*OK371-GAL4>UAS-myr::GFP*) at 79h APF following injection at 32h APF with a control solution and, (**D**), a solution containing TTX. (**E**) An axon terminal of a *VGlut* null motoneuron MARCM clone (green) alongside a *VGlut* heterozygous terminal in a filleted L3 larva stained with an antibody against VGlut (magenta). (**F**) The anterior-most arborisation in segment A3 (A3-A) of a heterozygous *VGlut* control expressing *myr::GFP* and *mCD8::GFP (VGluř^MJ×^-GAL4*) at 85h APF (adjacent arborisations have been removed on a slice by slice basis for clarity) alongside (**G**), an equivalent *VGlut* null MARCM clone at 80h APF (H&I) Arborisations of *VGlut* null A3-A MARCM clones did not differ significantly from controls in arbor area (controls: 44643 ± 2982*μ*m^2^, *VGlut*. 46518 ± 3083*μ*m^2^, n1,2 = 10, t(18) = 1.38, p = 0.18, t-test, two-tailed), but had marginally greater total arbor lengths (controls: 1812 ± 134*μ*m, *VGlut:* 2024 ± 216*μ*m, n1,2 = 10, t(18) = 2.64, p = 0.02, t-test, two-tailed). (**J**) Kymographs show changes in muscle GCaMP6m (*Mef2-GAL4*) Δf in response to motoneuron activation using the warmth-gated ion channel TRPA1 (*VGlut-LexA*) at 49h, 54h, 68h and 71h APF. Red bars indicate time at the restrictive temperature (22°C), green bars indicate time at the permissive temperature (30°C). (**K**) Images show muscle GCaMP6m Δfs before and after activation of motoneurons with dTRPA1 at 71h APF. (**L**) Time series showing the same arbor segment at 34h, 48h and 56h APF shows the shift in the organisation of BRP::RFP puncta from a predominance at branch points (white arrowheads) to a distribution along branch lengths (yellow arrowheads) (**M**) At stages between 72h and 77h APF, 12.55 ± 5.79% of total BRP::RFP puncta are found at branch points/bases of filopodia, although the majority (89.38 ± 6.73%) of branch points/bases of filopodia host puncta (n = 5). (**N**) Puncta of endogenous, GFP labelled BRP (indicated by arrowheads) are larger and more homogenous in diameter during early arbor growth (32h APF) than at later stages (70hAPF), when outgrowth has ceased (*OK371-GAL4>UAS-mCD8::Cherry)*. (**O**) Diameters of endogenous BRP::GFP puncta measured as the full width at half maximum of peaks in fluorescence are significantly greater at 32h APF (435.8 ± 177.8nm, n = 144) than at 72h APF (319.6 ± 87.9nm, n = 177) (Mann-Whitney U = 6935, p < 0.0001, two-tailed). (**P**) GFP tagged *GluRIIE* (FlyFos (Sarov *et al*., 2016)) driven under the control of the native transcriptional machinery shows an absence of Glutamate receptors from the postsynaptic membrane as late as 48h APF. (*OK371-GAL4>UAS-mCD8::Cherry)*. (**Q**) At 85h APF, conspicuous GluRIIE::GFP clusters (arrowheads) adorn the axon terminals. Bars represent SDs. Scale bars: 50*μ*m (**A,C,D,F,G**), 10*μ*m (**E**), 100*μ*m (**K**), 25*μ*m (**L**), 5*μ*m (**N,P,Q**).

To address directly whether arbor growth can take place independently of activity-evoked neurotransmission we silenced neurons by intravital injection of the sodium channel blocker tetrodotoxin (0.05mM TTX). Of the 8 animals injected between 24h and 32h APF, 6 showed no detectable calcium activity at 78h APF (Figure 3B) and 2 showed very weak transients restricted to small parts of branches. In contrast, buffer injected controls gave expected levels of presynaptic activity (n = 5). At 79h APF we found the arborisations of animals injected with TTX to be morphologically indistinguishable from those injected with buffer (Figures 3C&D).

In addition to evoked neurotransmission, spontaneous neurotransmitter release, which is responsible for miniature postsynaptic potentials (mEPSPs), has been shown to play a key role in the growth and development of neurons (Choi *et al.*, 2014; Andreae & Burrone, 2015). To determine if spontaneous neurotransmission is important for the growth of PM-Mn axonal arborisations we generated PM-Mn clones homozygous mutant for the vesicular glutamate transporter (*VGlut*) using the null allele *Df(2L)VGlut*^2^. This allele has been previously shown to completely block both evoked and spontaneous glutamatergic neurotransmission (Daniels *et al.*, 2006). Antibody staining confirmed that *Df(2L)VGlut*^2^ MARCM clones have a complete loss of VGlut protein in motoneurons (Figure 3E). To assess the effect of removing *VGlut* on arbor growth we performed morphometric analysis on clones of the anterior-most motoneurons of segment A3 (A3-A) in females staged between 80h and 90h APF (Figures 3F&G). No significant difference was found between the area of coverage of *VGlut* null A3-A arborisations and controls (Figure 3H), although *VGlut* nulls were marginally greater in total arborisation length (Figure 3I).

The timeline of neural activity and the *VGlut* null data strongly suggests that synaptic transmission does not play a role during the ‘elaborative phase’ of growth of the pleural neuromuscular system. To determine when synaptic transmission is ‘physiologically’ possible we artificially stimulated neurons using the warmth-gated ion channel TRPA1 (Hamada *et al.*, 2008), and measured postsynaptic calcium responses in the pleural muscles with GCaMP6m. Before 49h APF no calcium events were recorded in the muscles (n = 2) (Figure 3J). Between 50h and 59h APF a few large calcium events were recorded in the muscles but in each case these occurred either during the final seconds of stimulation or just after ramping down the temperature (n = 2). In contrast, when stimulating at stages between 68h and 72h APF we found robust, rapid and sustained postsynaptic calcium activity (Figures 3J&K). Between 68-72h APF calcium events occurred at a significantly greater rate at the permissive temperature (30°C) than at the restrictive temperature (22°C) (6.15 ± 5.60 minute^-1^ vs 0.93 ± 0.81 minute^-1^, Mann-Whitney U = 6.0, n = 7, p = 0.02, two-tailed). These data indicate that synaptic transmission does not take place within this system before 60h APF.

The lack of impact on growth from removing synaptic transmission made us look more closely at the organization of presynaptic machineries at different times during development. To do this we measured the distribution of BRP::RFP puncta in the same arborisation at 12 hour intervals from 34 to 58h APF (Figure 3L). As described previously, at 34h APF BRP::RFP puncta are found almost exclusively at branch points and at the bases of filopodia (white arrowheads). By 48h APF, in addition to the puncta at branch points, many puncta are also found along the lengths of branches, between branch nodes (yellow arrowheads). By 58h APF all but the most proximal branches are lined with large numbers of BRP::RFP puncta. Finally, an analysis of the distribution of puncta at stages 72-77h APF found that just 12.6 ± 5.8%, n = 5 arborisations of individuals) of puncta are at branch points, the rest being distributed along branch lengths (Figure 3M). The majority, 89.4 ± 6.7%), of branch points were still found to have puncta.

In addition to distribution, we used the BRP::GFP protein trap line to measure changes to BRP puncta size (Figures 3N&O). At early stages, BRP::GFP puncta diameters were found to be far more heterogenous, shown by the difference in standard deviation between the groups, found to be significant by an F-test of equality of variances (F_143,176_ = 4.092, p < 0.0001). We also found that BRP::GFP puncta became significantly smaller over the course of development (Figure 3O).

These smaller and more homogenously sized BRP puncta in later development highlighted that the puncta we see in early PM-Mn arborisations may not actually be synapses. To explore this idea we asked when glutamate receptors appear in the muscles. No detectable GluRIIE was evident at 48h APF (Figure 3P) yet by 85h APF clear clusters of GluRIIE were found apposed to the branches (Figure 3Q). Thus, the changes in the organisation of synaptic machineries over the course of development suggest that the early accumulations of presynaptic components do not represent differentiated synapses.

### A role for Neuroligin1-Neurexin in PM-Mn arborisation growth

Neurexin-Neuroligin signalling is capable of driving synapse formation in many contexts and in the *Xenopus* tectum regulates arbor growth in an activity dependent manner (Haas *et al*, 2010). Though our data ruled out activity playing a key role in PM-Mn axonal arbor growth, we next sought to test whether these synaptic adhesion proteins are important.

We generated homozygous mutants using the null alleles, *Nlg1*^*e*×2.3^, Nlg1^1960^ (Banovic *et al*., 2010) and Nrx^241^ (Li *et al*., 2007) in combination with chromosomal deficiencies. These combinations are viable into adulthood. The arborisations of both *Nlg1* and *Nrx* mutants showed comparable defects with large reductions in coverage relative to wild-type controls (Figures 4A&B).

**Figure 4.**
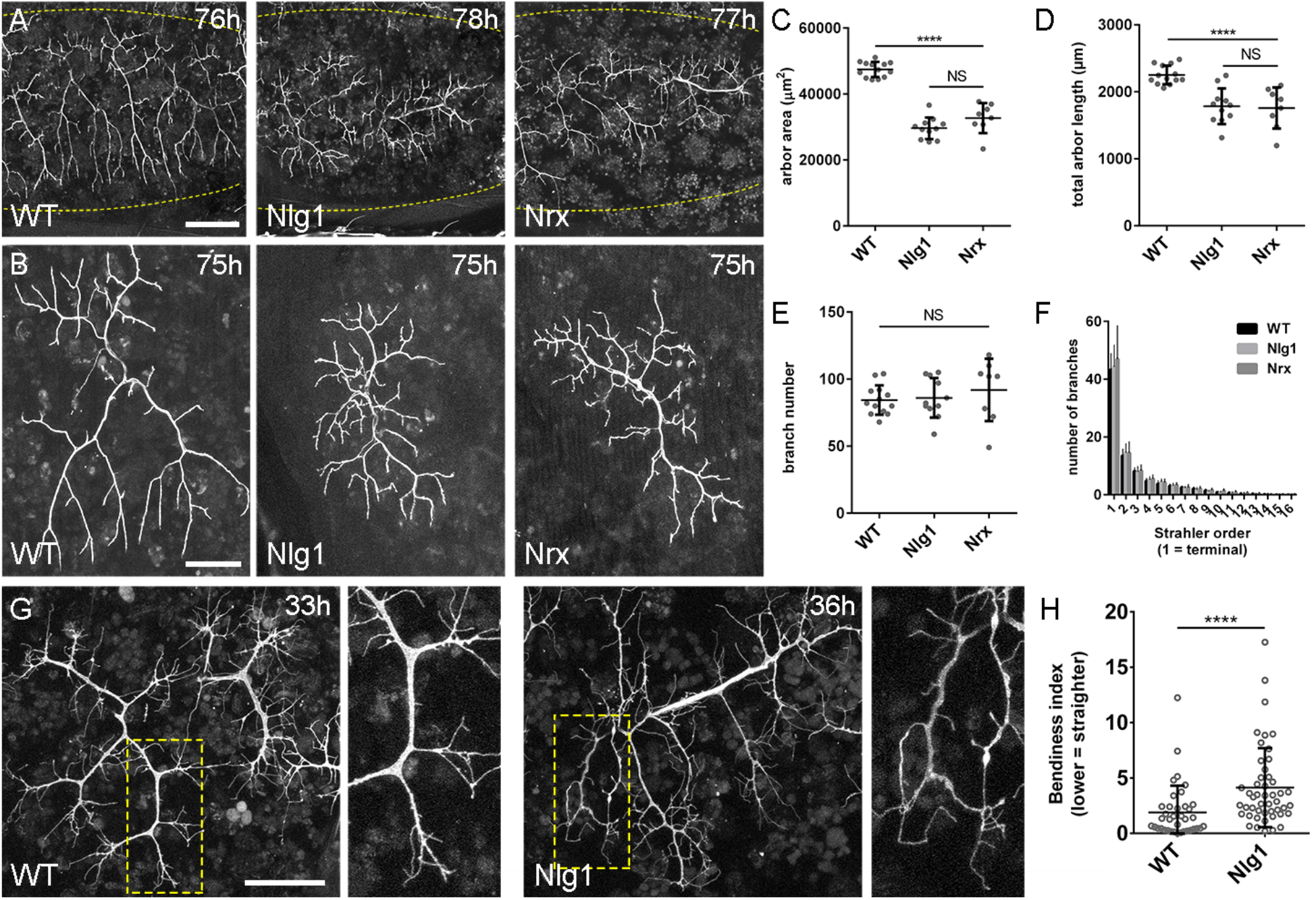
*Neurexin* and *Neuroligin 1* are required during early arbor growth. **(A)** Arborisations in segments A3-A5 in a wild-type control, an *Nlg1* null and an *Nrx* null staged between 76h and 78h APF (VGlut-LexA>myr::GFP). (**B**) Anterior-most arborisations in segment A3 (A3-A) at stages between 75h and 85h APF. Arborisations belonging to other motoneurons have been removed using image processing. (**C-F**) Morphometric analysis of A3-A arborisations of *Nlg1* nulls (n = 11), *Nrx* nulls (n = 8) and wild-type controls (n = 13) staged from 75h to 85h APF. *Nlg1* and *Nrx* nulls had significantly lower (**C**), arbor areas (controls: 47445 ± 2298*μ*m^2^, *Nlg1:* 29616 ± 3259*μ*m^2^, *Nrx:* 32696 ± 4565*μ*m^2^. *Nlg1* vs controls: t(22) = 15.67, p < 0.0001. *Nrx* vs controls: t(19) = 9.89, p < 0.0001. t-tests, two-tailed) and (**D**), total arbor lengths (controls: 2251 ± 140*μ*m, *Nlg1:* 1785 ± 265*μ*m, *Nrx:* 1758 ± 306*μ*m. *Nlg1* vs controls: t(22) = 5.51, p < 0.0001. *Nrx* vs controls: t(19) = 5.07, p < 0.0001. t-tests, two-tailed), but not (**E**), total branch numbers (controls: 84.38 ± 10.97, *Nlg1:* 86.00 ± 14.74, *Nrx:* 91.88 ± 23.27. *Nlg1* vs controls: t(22) = 0.31, p = 0.76. *Nrx* vs controls: t(19) = 1.00, p = 0.33. t-tests, two-tailed). (**F**) *Nlg1* nulls, *Nrx* nulls and controls were no different in their topological organisation. Nulls did not differ from controls in their total number of highest order (terminal) branches (controls: 43.31 ± 5.59, *Nlg1:* 44.27 ± 7.56, *Nrx:* 47.00 ± 11.50. *Nlg1* vs controls: t(22) = 0.36, p = 0.72. *Nrx* vs controls: t(19) = 0.99, p = 0.33. t-tests, two-tailed) or in their total number of orders (controls: 11.62 ± 1.94, *Nlg1:* 11.27 ± 2.15, *Nrx:* 11.88 ± 2.36. *Nlg1* vs controls: t(22) = 0.41, p = 0.67. *Nrx* vs controls: t(19) = 0.27, p = 0.79. t-tests, two-tailed). (**G**) At early stages of growth *Nlg1* null arborisations bear branches which are precociously long, yet more tortuous and with fewer side branches than wild-types. (**H**) Branch Bendiness measured as the percentage difference between the actual length and the straight-line length of primary (terminal) and secondary branches. Branches of *Nlg1* nulls are significantly less straight (4.13 ± 3.57%, n = 48) than controls (1.89 ± 2.41%, n = 38) at stages between 32h and 36h APF (Mann-Whitney U = 441, p < 0.0001, two-tailed). Bars represent SDs. Scale bars: 100*μ*m (**A**), 50*μ*m (**B,G**).

Morphometric analysis showed areas of coverage of *Nlg1* and *Nrx* null arborisations were significantly lower than wild-type controls (Figure 4C), as were total arbor lengths (Figure 4D). Interestingly however, there was no difference in the total branch numbers (Figure 4E). *Nrx* and *Nlg1* null arborisations were not significantly different from each other in any of these measurements. To determine if arbor complexity was different in the nulls we used the Strahler method of branch ordering (Figure 4F). Firstly, the groups showed no differences in their number of branch orders. Secondly, the total number of lowest order (terminal) branches was not significantly different between conditions; a trend which continued for subsequent orders, with each group showing remarkably similar numbers of branches at every level.

The morphology of late stage *Nlg1* and *Nrx* nulls points toward a requirement for these proteins but does not tell us *when* they are required. To address the timing of requirement, we compared *Nlg1* null arborisations with those of wild-type controls at 30-36h APF. PM-Mns in *Nlg1* nulls at this stage generate similar numbers of very dynamic filopodia compared to controls. A clear difference however was the far greater ‘bendiness’ or the tortuosity of *Nlg1* null branches (Figure 4G). To evaluate this, we scored primary and secondary branches of *Nlg1* null and control arborisations staged between 30h and 36h APF using an index of bendiness (Figure 4H). This was calculated from the percentage difference between the actual length of branches and the straight-line distance between their nodes. *Nlg1* null branches were found to be significantly less straight than those of the controls (Mann-Whitney U = 441, p < 0.0001, twotailed). This early phenotype indicates a requirement for Nlg1 signalling during the very initial phases of pleural neuromuscular development.

Although the full mutants showed robust and consistent phenotypes, there is limitation on what these can tell us. For example, they cannot reveal to us whether the phenotype is due to a failure of ‘local interactions’ between the developing synaptic partners. To address this we generated FLEX-based, FlpStop tools (Fisher *et al*, 2017) that allow conditional disruption of endogenous Nlg1 expression in a clonal fashion (Figures 5A&B).

**Figure 5.**
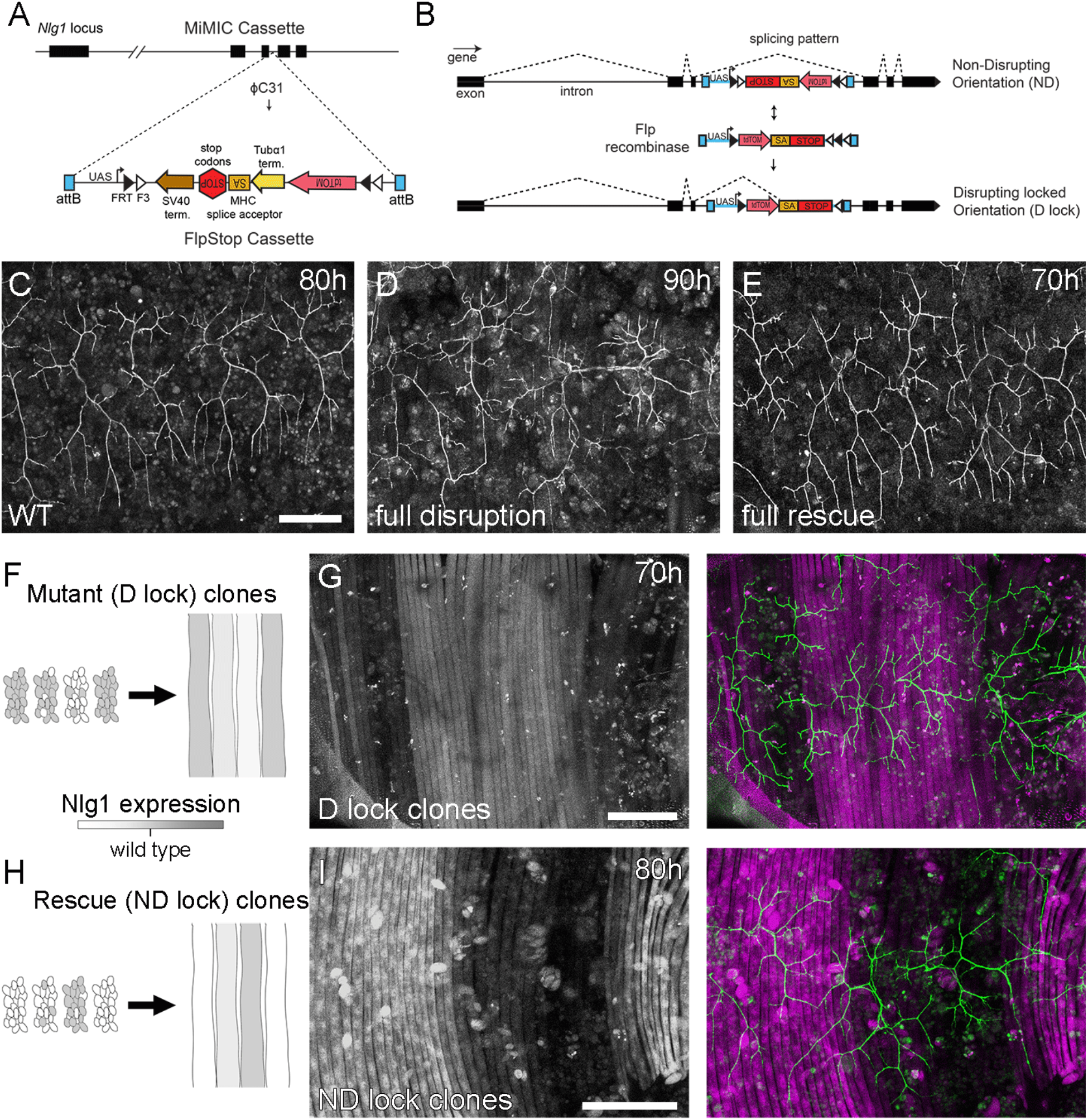
Mosaic analysis shows a local role for endogenous *Neuroligin 1* during arbor growth. (**A**) A schematic shows the directed insertion of the FlpStop construct into the 3^rd^ coding intron of the *Nlg1* genomic locus by recombinase mediated cassette exchange (RMCE) with a MiMIC integration site. (**B**) A schematic shows the mechanism of FlpStop action for the conditional disruption of endogenous *Nlg1*. In the initial non-disrupting (ND) orientation the splice acceptor and stop codons are on the non-coding strand and thus do not effect gene function. Upon inversion by Flp recombinase the splice acceptor and stop codons are brought into frame on the coding strand, resulting in disrupted gene expression. The cassette is locked into this disrupting (D-lock) orientation by a FLEx switch (Schnütgen *et al*., 2003). In addition, inversion brings the coding sequence for tdTomato into proximity with a UAS sequence, enabling GAL4 driven expression. FlpStop lines were also generated in the initially disrupting (D) orientation, which allows rescue of *Nlg1* (ND-lock clones) and ‘turns on’ tdTomato (Schematic not shown). (**C**) Arborisations at 80h APF in a wild-type control (*VGlut-LexA>myr::GFP)*. (**D**) Complete induction of *Nlg1* D-lock (disrupting) clones using *hsFlp^122^* and a long heat-shock produces arbor growth defects comparable to *Nlg1* mutants. (**E**) Germline ND-lock rescues arbor growth to a near wild-type phenotype. (**F**) Model shows the formation of *Nlg1* deficient fibres from the fusion of D-lock clonal myoblasts induced using *hsFlp^122^* and a short heat shock at larval L3 stage. (**G**) Arbor growth (*VGlut-LexA*) onto regions of D-lock muscle clones (*Mef2-GAL4*) is disrupted, whereas arbor growth on non-clonal regions is close to wild-type. (**H**) Model shows the rescue of *Nlg1* expression in clonal fibres formed from the fusion of ND-lock clonal myoblasts induced using *hsFlp^122^* and a short heat shock at larval L3 stage. (**I**) Arbor growth onto ND-lock muscle clones is close to wild-type, whereas growth on non-clonal regions is disrupted. Scale bars: 100*μ*m (**C,D,E,G,I**).

Using a MiMIC insertion within the 3^rd^ coding intron of *Nlg1* we generated lines capable of either rescuing *Nlg1* in a mutant background or mosaically disrupting *Nlg1* in a wild-type background. For complete gene disruption, FlpStop lines were used in a heterozygous condition with a deficiency covering *Nlg1*. To test the ability of the non-disrupting orientation (ND) to disrupt Nlg1 expression upon cassette inversion, *hsFlp^122^* was used to induce large numbers of D-lock clones. At 90h APF, a very similar phenotype was observed to *Nlg1* nulls (Figure 5D). To test the ability of the disrupting orientation (D) to be converted into a non-disrupting allele (ND-lock), a germ-line inverted stock was generated. In this case, arbor growth was rescued to a near wild-type phenotype (Figure 5E).

To test the local requirement of Nlg1 we induced small numbers of FlpStop muscle precursor clones that generated fibres containing varying numbers of *Nlg1+* nuclei (Figures 5F&H). Looking at stages between 70h and 85h APF, the organisation of arborisations, relative to the pattern of clonal muscle fibres, suggest a local role for Nlg1 mediated adhesion in branch growth. In the initially non-disrupting orientation, terminals growing on non-clonal/low level clonal muscle appear to be phenotypically closer to wild-type (Figure 5G). In contrast, branches belonging to neurons which had grown on muscle with more clones, displayed phenotypes comparable to complete Nlg1 nulls (see Figure 4B). Using the disrupting orientation allele the reverse situation was observed (Figure 5I). Branches growing across clones in which Nlg1 expression was rescued displayed more wild-type morphologies than those growing on non-clonal fibres (n = 8 muscle mosaics). This is demonstrated in Figure 5I by the two arborisations spanning clonal and non-clonal muscles; segments on the wild-type clonal regions show extensive growth, whereas branches on the non-clonal regions show a reduction in growth comparable to complete nulls (n = 7 muscle mosaics).

### Dynamic complexes of ‘synaptic’ adhesion proteins stabilise filopodia and drive branch growth

One prediction from these mutant data is that Nlg1 could ‘prepattern’ PM neuromuscular junctions, as is seen with acetylcholine receptors in zebrafish somatic muscles (Panzer *et al.*, 2006). To test this idea we looked at the location of postsynaptic Nlg1 with a GFP tagged version of Nlg1 (Banovic *et al*., 2010). At 35h APF we found that Nlg1::GFP faintly labels the entire postsynaptic membrane, but forms strong puncta only at sites that directly appose the axon terminals (Figure 6A). To understand what was happening we simultaneously looked at the dynamics of the growing neurons and the localisation of Nlg1. Higher magnification imaging showed Nlg1 puncta were found concentrated on growing branches, particularly at sites of filopodia growth, along the lengths and also at the very tips of filopodia (Figure 6B and movie 7).

**Figure 6.**
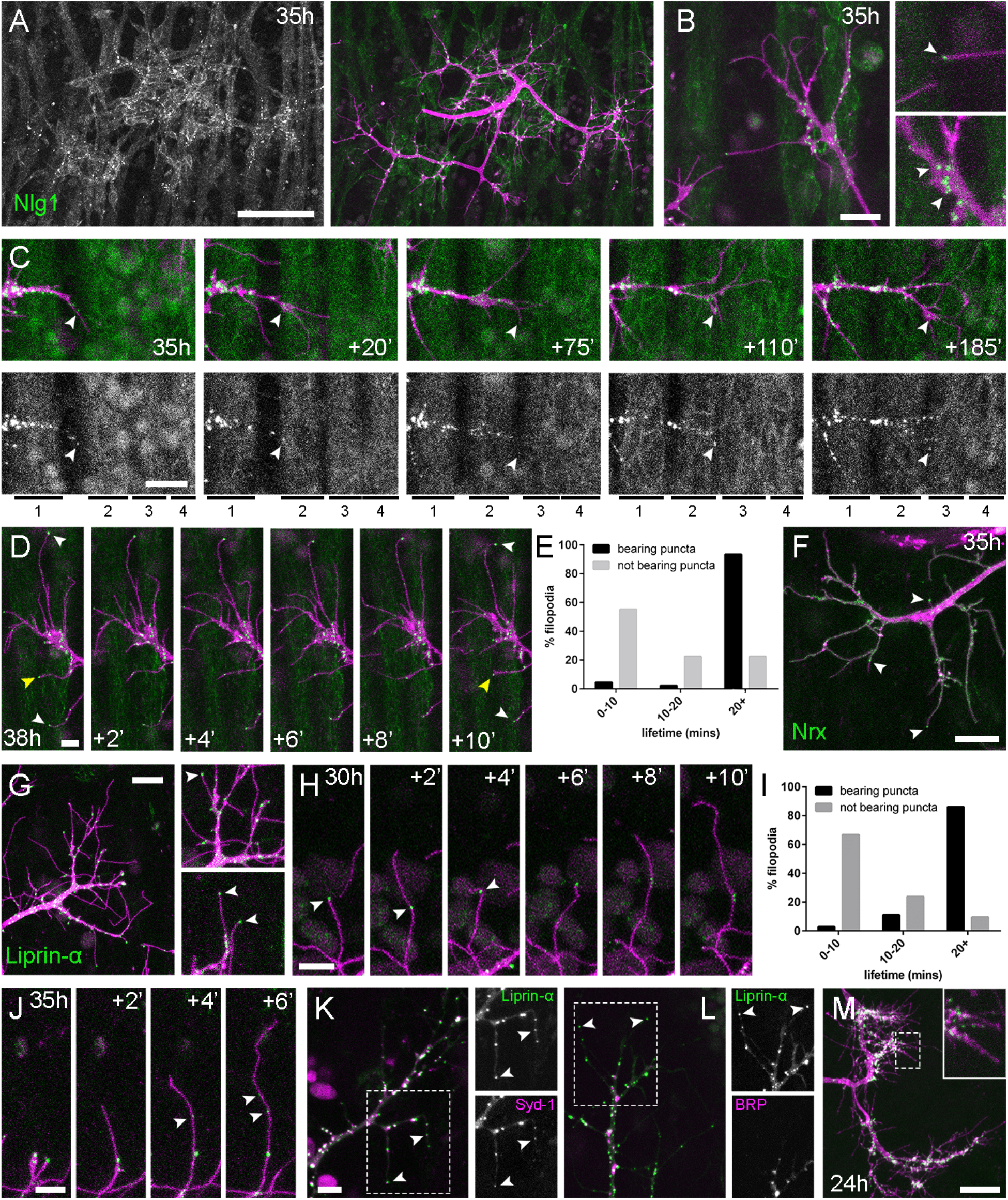
Neuritic Adhesion Complexes (NACs) incorporating *Neurexin* and *Neuroligin 1* stabilise filopodia during arbor growth. **(A)** At 35h APF *Nlg1::GFP* expressed in muscles (*Mef2-GAL4*) (green) formed puncta which arranged exclusively in apposition with the axon terminals on the postsynaptic membrane (*VGlut-LexA>myr::tdTomato*) (magenta). (**B**) Higher magnifications show the concentration of Nlg1::GFP puncta at sites of branch growth and their apposition to filopodial tips. (**C**) Time series shows the extension of a branch across a muscle field at 35h APF. Muscles are numbered from anterior to posterior. Rounds of filopodia extension and stabilisation are concomitant with the recruitment of Nlg1::GFP puncta (indicated by white arrowheads) to their tips. (**D**) Time series shows the relationship between the dynamics of filopodia and Nlg1::GFP puncta. Puncta indicated by white arrowheads mark the tips of filopodia which persist. Yellow arrowhead indicates a punctum which marks the limit of filopodial retraction (**E**) Histogram showing lifetimes of filopodia with Nlg1::GFP puncta at their tips (n = 45) and without puncta at their tips (n = 49). Filopodia bearing Nlg1::GFP puncta were significantly longer lived than those without puncta (with puncta: 19.36 ± 2.82 mins, without puncta: 10.73 ± 6.88 mins, Mann-Whitney U = 305.5, p < 0.0001, two-tailed). (**F**) *Nrx::GFP* (green) expressed in an arborisation with *mCD8::Cherry* (magenta) at 35h APF (*OK371-GAL4).* Arrowheads indicate puncta positioned on filopodia. (**G**) *Liprin-a::GFP* (green) expressed in an arborisation with *mCD8::Cherry* (magenta) at 35h APF (*OK371-GAL4)*. Cutaway images show Liprin-α::GFP puncta in branches and at the tips of filopodia (arrowheads). (**H**) Time series shows the extension of a filopodium and its retraction back only as far as a Liprin-α::GFP punctum (arrowheads). (**I**) Histogram showing lifetimes of filopodia with Liprin-α::GFP puncta at their tips (n = 36) and without puncta at their tips (n = 21). Filopodia bearing Liprin-α::GFP puncta were significantly longer lived than those without puncta (with puncta: 19.44 ± 1.96 mins, without puncta: 8.81 ± 5.88 mins, Mann-Whitney U = 51.5, p < 0.0001, two-tailed). (**J**) Time series shows the rapid precipitation of new Liprin-α::GFP puncta (indicated by arrowheads) within a filopodium. (**K**) *Liprin-a::GFP* and *Syd-1::Straw* co-expressed with *OK371-GAL4* form puncta in growing axon terminals which are highly coincident, including at filopodia tips, as indicated by arrowheads. The slight misalignment of some puncta is a result of the latency in imaging channels sequentially. (**L**) In contrast, Liprin-α::GFP and BRP::RFP puncta are coincident only at a few sites along branch lengths and at branch points. Unlike Liprin-α::GFP puncta (arrowheads), BRP::RFP puncta never stabilise within filopodia. (**M**) *Liprin-α::GFP* expressed in a central interneuron (*Rn2-FLPout*) forms puncta in growing axon terminals, including on filopodia at 24h APF. Bars represent SDs. Scale bars: 50*μ*m (**A**), 10*μ*m (**B,F,G**), 20*μ*m (**C,M**), 5*μ*m (**D,H,J**).

To look at the timing of Nlg1 recruitment with regards to branch growth we imaged at 5-minute intervals from 35h APF (Figure 6C). From these time lapses we found that Nlg1::GFP puncta are recruited directly onto filopodia and the tips of growing branches. As a result, branch growth occurs as a highly coordinated sequence whereby the arrival of Nlg1::GFP on exploratory processes precedes their stabilisation and maturation into stable branches in an iterative manner.

The dynamic recruitment of Nlg1::GFP puncta to growing branches and filopodia suggests a significant role for Nlg1 during the earliest phases of arbor construction. To look for a possible relationship between Nlg1::GFP puncta and filopodia/branch dynamics, we took faster time-lapses with 1-minute intervals of arborisations between 30 and 35h APF. Shown in the series in Figure 6D, we found that Nlg1::GFP puncta in apposition to growth cones and filopodia are often stable for many minutes. What is more, these puncta at the tips of filopodia (white arrowheads) appeared to correlate with filopodia longevity. In addition, we found that puncta mark the limits of filopodial retraction (yellow arrowhead). To assess this relationship, we compared the lifetimes of filopodia with and without Nlg1::puncta in time-lapses of 7 individuals. Due to photobleaching, these sequences were limited to ~ 20 minutes. Very few incidents were observed in which newly generated filopodia recruit an Nlg1::GFP punctum. Therefore, instead of analysing newly generated filopodia, only filopodia in existence at the beginning of recording were considered in the analysis. The graph in Figure 6E shows that 55.1% of filopodia not apposed to puncta (n = 49 filopodia) were lost within 10 minutes, 22.5% were lost within 20 minutes and a further 22.5% lasted longer than the duration of the movies. On the other hand, only 4.4% (2 filopodia) of the population apposed to puncta (n = 45 filopodia) were lost within 10 minutes, 2.2% (1 filopodium) were lost within 20 minutes and 93.3% survived for longer than the duration of the recordings. Statistical analysis found that filopodia tipped by Nlg1::GFP puncta are significantly longer lived than filopodia not bearing puncta (Figure 6E).

The major trans-synaptic binding partners of Neuroligins are the Neurexins. Thus, a natural prediction would be that presynaptic Nrx should mirror the postsynaptic distribution of Nlg1::GFP. We expressed GFP tagged *Nrx1* (hereon termed *Nrx*) in the PM-Mns and found that Nrx::GFP (Banovic *et al*., 2010) forms puncta during early arbor growth that localise to points of filopodial growth and along filopodial lengths, including tips of filopodia (Figure 6F and movie 8). The low signal and rapid bleaching of Nrx::GFP made it difficult to follow Nrx dynamics *in vivo* over longer periods.

Alongside Nrx we also looked at two key players in presynaptic development; Syd-1 and Liprin-α. Liprin-α (Lar interacting protein) is a scaffolding protein that is one of the first components recruited to trans-synaptic Nrx-Nlg complexes (Owald *et al*, 2012). *Liprin-α::GFP* (Fouquet *et al*., 2009) was expressed in motoneurons with *OK371-GAL4*. Much like our Nrx::GFP data, Liprin-α::GFP forms distinct puncta at branch terminals and within filopodia (Figure 6G). Similarly to postsynaptic Nlg1::GFP, we found that presynaptic Liprin-α::GFP puncta regularly marked the limits of filopodial retraction (Figure 6H) and appeared to correlate with filopodial stability (see movie 9). Indeed, filopodia bearing Liprinα::GFP puncta were significantly longer lived than those not (Figure 6I). It appears that Liprin-α::GFP coalesces into puncta directly on filopodia (Figure 6J). To ask if Liprin-α::GFP puncta mark adhesion complexes we looked at the localisation of another known interactor, *Syd1. Syd1::GFP* (Owald *et al*., 2010), like *Liprin-α::GFP*, forms puncta which localise within growing branch terminals, including at the tips of filopodia (data not shown). To see if these 2 proteins colocalise to the same sites we expressed *Liprin-α::GFP* together with Strawberry tagged *Syd1* (Syd1::Straw) (Owald *et al*., 2010). The large majority of puncta of each protein were coincident, including those at the tips of filopodia (Figure 6K). In contrast, when *Liprin-α::GFP* and *BRP::RFP* were expressed together there was very little co-localisation at branch points and never at filopodia tips (Figure 6L).

Finally, to determine the subcellular distribution of Liprin-α::GFP in the central nervous system we looked at *Eve+* interneurons. Like in growing axon terminals of PM-Mn neurons, Liprin-α::GFP is localised to the tips and along the lengths of filopodia on the growing axonal output arborisations (Figure 6M).

### Nlg1-based adhesion complexes can direct a tropic mode of growth

The relationship between Nlg1::GFP puncta and filopodial dynamics suggests that it plays a role in the growth of axonal arborisations by providing adhesive stability to branches and filopodia during the early stages of growth. A direct prediction from this would be that if we manipulated the postsynaptic levels of Nlg1 at early stages we would see changes in PM-Mn growth. To explore this, we used *UAS-Nlg^untagged^* which is known to be expressed at high levels and cause a strong phenotype in larvae (Banovic *et al*., 2010).

At 33-35h APF, elevated postsynaptic Nlg1 levels resulted in compacted axon terminals with distal branches transformed into flattened growth cones with many filopodia (Figures 7A&B). This irrevocable effect on growth led to late stage arborisations with reduced branch length, territory and complexity (Figures 7C&D). These branches also maintained greater than usual numbers of filopodia into the later stages of development.

**Figure 7.**
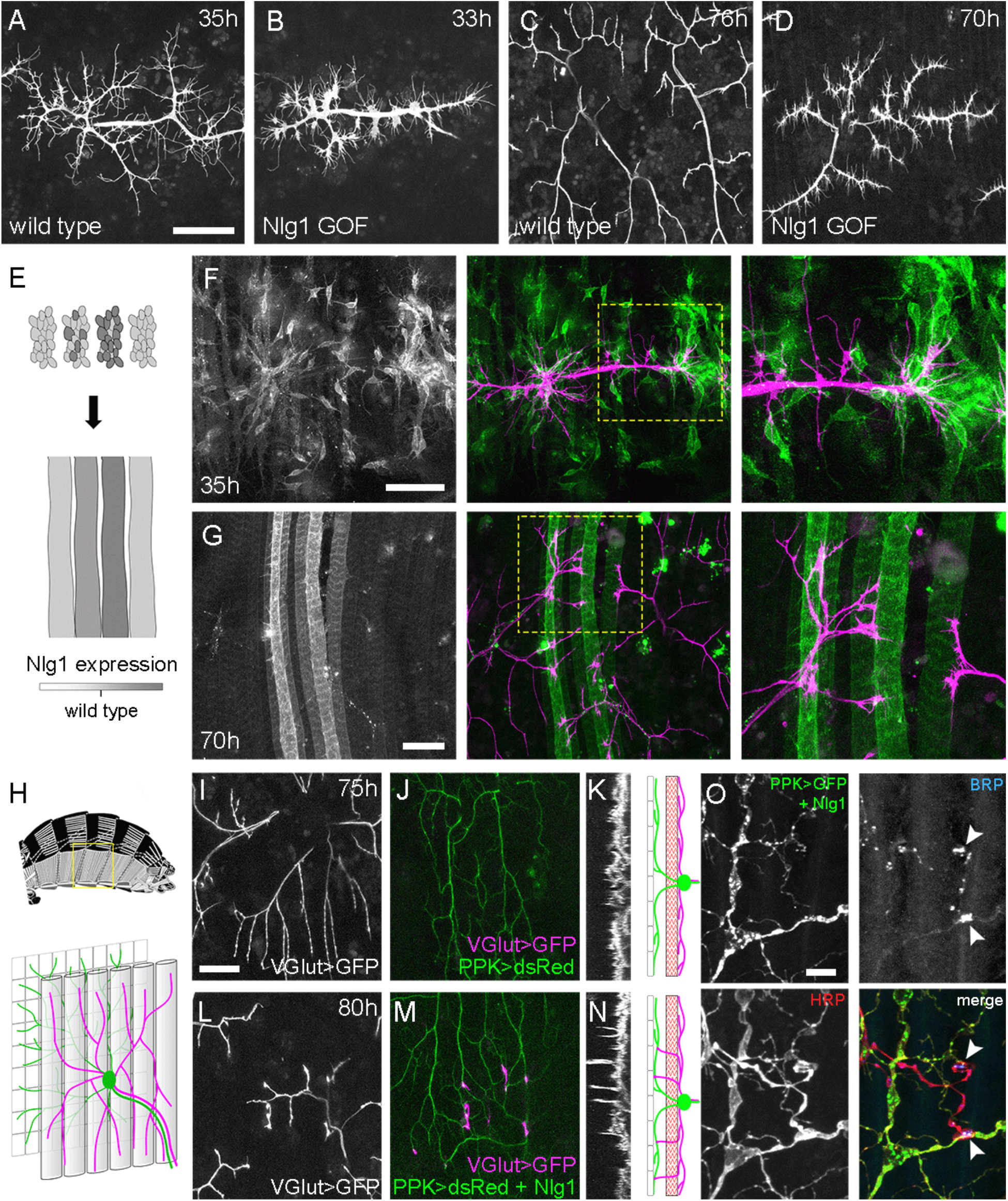
*Neuroligin 1* acts locally to drive early arbor growth. **(A-D)** Postsynaptic *Nlg1* overexpression severely impacts axon arbor growth. (**A**) A pair of arborisations at 33h APF expressing *myr::GFP (VGlut-LexA*) in a background of postsynaptic *Nlg1^untagged^* expression (*Mef2-GAL4)*. (**B**) Pair of control arborisations at 35h APF. (**C**) Arborisations at 70h APF in a background of postsynaptic *Nlg1^untagged^* expression. (**D**) Control arborisations at 76h APF. (**E**) Model of ‘flip-out’ *Nlg1^untagged^* and *myr::tdTomato* expressing muscle clone induction using *αTub84B-FRT.GAL80*. Muscle precursor clones induced with *hsFlp^122^* at larval L3 stages produce clusters of GAL80 negative myoblasts which fuse to form ‘stripes’ of clonal muscle fibres. (**F**) At 35h APF motoneuron branches (magenta) show preferential elaboration onto clonal *Nlg1^untagged^* expressing myotubes and myoblast clusters (green). (**G**) At 70h APF only axon branches in contact with clonal, *Nlg1^untagged^* expressing muscle fibres show a hyperstabilisation phenotype. Growth of these branches appears to have been directed preferentially along these fibres. (**H-O**) Ectopic expression of Nlg1 in class IV da sensory neurons drives changes in motoneuron axon arbor morphology. (**H**) A model of the pleural body wall shows the relative positions of class IV v’ada sensory input arborisations (green), pleural muscles (grey tubes) and motor axon arborisations (magenta). (**I**) Motoneuron axon terminals expressing *myr::GFP (VGlut-LexA*) at 80h APF in a wild-type control (**J**) the v’ada input arborisations in the same region expressing *dsRed (PPK-Gal4* (Grueber *et al*., 2003)) (**K**) a transverse projection shows the restriction of the motor axon arborisations to a single plane. This is represented in a transverse view of the pleural region which shows the separation of the v’ada arborisation (green) from the motor axon terminals (magenta) by the pleural muscles (red). (**L-N**) *Nlg1^untagged^* expression in class IV da sensory neurons causes motor axon branches to penetrate gaps between the muscle fibres to make contacts with the sensory arborisations. These aberrant branches are shown in the transverse view and run perpendicularly to the rest of the arborisation. (**O**) Abdominal fillet of a newly eclosed adult expressing *Nlg1^untagged^* and *CD8::GFP* in the class IV da sensory neurons. Anti-HRP reveals the motor axon terminals. BRP immunoreactivity reveals presynaptic specialisations at contacts between motor axons and the sensory arborisation (arrowheads). Scale bars: 50*μ*m (**A,B,C,D,F,G**), 25*μ*m (**I,J,L,M**), 5*μ*m (**O**).

Although full postsynaptic overexpression of *Nlg1^untagged^* revealed that axon terminals are sensitive to Nlg1 levels from very early stages of growth, to explore how different levels of Nlg1 signalling impacts branch growth at a local level we developed a clonal technique that generates patterns of expression much like a ‘Bonhoeffer stripe’ assay (Walter *et al*., 1987). With this ‘stripe assay’ we can present growing motoneurons with different levels of Nlg1^untagged^ to grow upon (Figure 7E). At early stages (35h APF), PM-Mn growth was directed onto myoblast clusters/developing myotubes that strongly expressed the *Nlg1^untagged^* (Figure 7F). By later stages (70h APF), branches in contact with these strongly expressing clonal fibres displayed the same hyper-stabilisation phenotype seen with full muscle expression. In contrast, branches from the neuron in contact with low to non-expressing fibres grew as normal, demonstrating unequivocally that Nlg1 impacts branch growth via local mechanisms (Figure 7G). Although branches contacting highly expressing fibres had reduced growth and branching, they could sometimes be seen to elaborate along a clonal fibre, suggesting a ‘tropic’ mode of growth is at play.

To further investigate Nlg1 signalling on branch growth we took advantage of the organisation of the peripheral nervous system in the abdominal body wall. As shown in Figure 7H, the class IV dendritic arborisation sensory neuron v’ada elaborates on the internal side of the epidermal cells. The peripheral neurites of v’ada are separated from the motor axon branches by the pleural muscles, which in turn receive innervation from the PM-Mns onto their internal surface. During early pupal development, motor and sensory arborisations are in very close proximity. We predicted that if we expressed Nlg1 in these usually ‘asynaptic’ sensory neurites the signalling would direct the growth of the PM-Mn arborisations into a novel territory. We expressed *Nlg1^untagged^* in v’ada neurons and then visualised the anatomy of the motoneurons. In controls branch growth was restricted exclusively to the inner surface of the muscle (Figures 7I&J). As highlighted by the transverse projection there is a relatively uniform layer of motoneuron terminals (Figure 7K). In contrast, when v’ada was made to misexpress *Nlg1^untagged^*, PM-Mn axon branches grow perpendicular to the normal arborisation between muscle fibres to contact v’ada sensory neurites (Figures 7L-N). In several cases further branching from these contacts increased the complexity of these ectopic PM-Mn branches.

One postulate of Vaughn’s tropic model of arborisation growth was that the stabilisation of contacts between synaptic partners would ultimately lead to functional connectivity (Vaughn, 1988). For this to be satisfied it would be expected that branches generated by such a mechanism would harbour mature synaptic terminals later in development. To test this in branches sculpted by ectopic *Nlg1^untagged^*, abdominal fillets were stained for the active zone marker BRP using the antibody NC82 and with the neural marker, anti-HRP. As shown in Figure 7O, conspicuous concentrations of BRP were found at each contact. Alongside this, we also found strong VGlut immunoreactivity at these ectopic synaptic contacts (data not shown).

### Physiological consequences of Nlg1 disruption

We have shown a very early requirement for Nlg1 for establishing branch structure. The ectopic contacts that PM-Mns make on v’ada neurites appear to mature into hemisynapses. These data show that following contact and stabilization there is a hierarchy of events that ultimately leads to synapse formation. To explore if Nlg1-Nrx disruptions impact synapse formation we stained against BRP to assess the density of active zones within the branches of *Nlg1* nulls, *Nrx* nulls and animals expressing *Nlg^untagged^* in muscles at pupal stages close to eclosion (Figure 8A). Boutons in all these animals were considered as engorgements containing concentrations of synaptic puncta. We found that the density of synapses was not significantly different between *Nlg1* nulls or *Nrx* nulls and wild-type controls, although it was slightly higher in *Nlg1* gain-of-function animals (Figure 8B). This suggests that global aspects of active zone development may be largely unaffected by loss of *Nlg1* or *Nrx*. One might expect that functional deficits arising from a failure in appropriate Nlg1 signalling could be due to either defective morphology and to changes in the transmission or both. To explore this, we looked at *GCaMP6m* expressed in the muscles as a proxy for measuring muscle depolarisation at late pupal/pharate stages just prior to eclosion. In wild-type controls, muscle calcium events at this stage usually occur in synchrony across the muscle field (Figures 8C&D and movie 10). In contrast, in *Nlg1* gain-of-function pupae, calcium events were more common in some groups of fibres than others, resulting in far less synchronicity (Figures 8E&F and movie 11).

**Figure 8.**
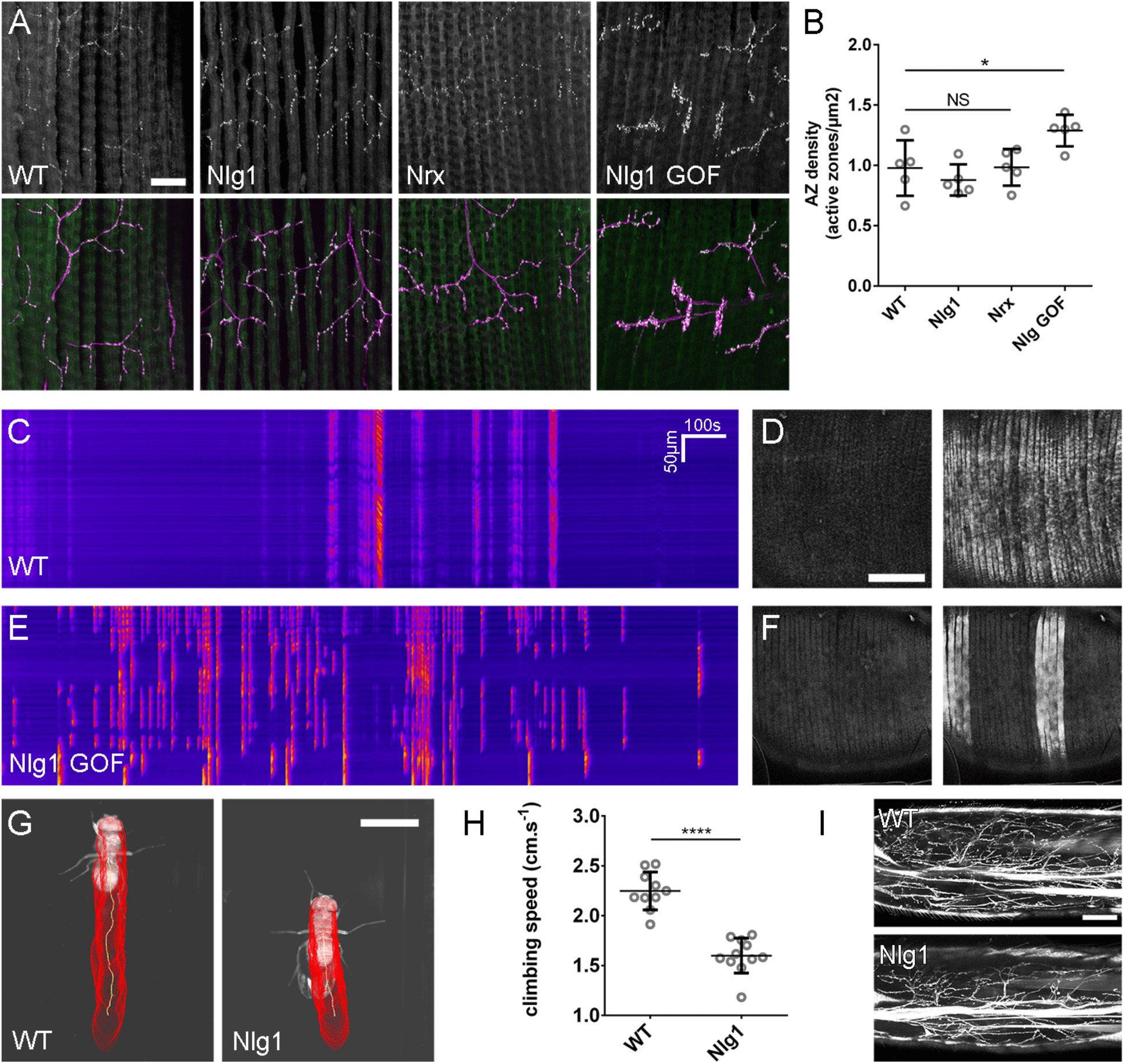
The roles of *Nlg* and *Nrx* in NAC dependent arbor construction are decoupled from synapse formation and neurotransmission. **(A-B)** Synapse formation is unaffected by disruptions to Nrx-Nlg signalling. (**A**) Antibody staining against BRP reveals active zones at the pleural neuromuscular junctions in adult abdominal fillets from a wild-type control, an *Nlg1* null, an *Nrx* null and an *Nlg1* gain-of-function (*Mef2-GAL4>Nlg1-untagged)*. (**B**) Active zone densities calculated from BRP puncta in synaptic terminals. No significant differences were found between the densities of active zones in controls (0.98 ± 0.23*μ*m^-2^, n = 5) and *Nlg1* nulls (0.88 ± 0.13*μ*m^-2^, n = 5) or controls and *Nrx* nulls (0.98 ± 0.15*μ*m^-2^, n = 5) (*Nlg1* vs controls: Mann-Whitney U = 9, p = 0.53, two-tailed. *Nrx* vs controls: Mann-Whitney U = 12, p = 0.94, two-tailed). A small, but significant difference was found between active zone densities in *Nlg1* gain-of-function terminals (1.29 ± 0.13*μ*m^-2^, n = 5) and controls (Mann-Whitney U = 1, p = 0.0159, twotailed). (**C-F**) *Nlg1* overexpression causes changes to arbor morphology which result in altered connectivity and transmission. Kymographs show muscle GCaMP6m Δf in a control, (**C**), and in a pupa also expressing *Nlg1^untagged^* in muscles (*Mef2-GAL4*) staged within 10 hours of eclosion, (**D**). In controls, calcium events are largely synchronous across the entire muscle field whereas in gain-of-function pupae events are often restricted to muscle subsets. This is demonstrated by the breaks in the light-coloured bars on the gain-of-function kymograph as well as in the sequential images (**D&F**). (**G-I**) Loss of *Nlg1* has a pronounced impact on motor ability (**G**) Automated body tracking performed using FlyLimbTracker (Uhlmann *et al*., 2017) in Icy (de Chaumont *et al*., 2012) of a control and an *Nlg1* null, 2-days post-eclosion (**H**) Control flies have a significantly faster climb speed than *Nlg1* nulls (controls: 2.25 ± 0.19 cm.s^-1^, n = 10. *Nlg1:* 1.60 ± 0.18 cm.s^-1^, n = 11. t(8.15) = 19, p < 0.0001, t-test, two-tailed). (**I**) Motoneuron axon terminals (*VGlut-LexA>myr::GFP*) innervating the femur 1-day post-eclosion of an *Nlg1* null and of a control. Bars represent SDs. Scale bars: 20*μ*m (**A**), 100*μ*m (D,F,I), 0.2cm (**G**).

To determine the functional consequence of disruption to Nlg1-Nrx signalling, we also assessed locomotor ability of *Nlg1* null flies in a climbing assay (Figure 8G). Using videography and an automated tracking software we found that the climbing speed of *Nlg1* nulls (n = 11) was significantly lower than controls (n = 10) (Figure 8H) (t(8.15) = 19, p < 0.0001, t-test, two-tailed). We found the leg motoneurons disrupted in *Nlg1* nulls, suggesting that some of this deficit may be the result of either disrupted growth of leg motoneurons arborisations or in central circuits (Figure 8I).

## Discussion

### PM-Mn axonal arborisations use a dynamic ‘synaptotropic-like’ mode of growth

Building arborisations of the right size and shape is critical for proper neural circuit development and function. Live imaging studies in vertebrate brains show that neuronal growth is highly dynamic and that structures which appear to be *nascent synapses* play a key role in the development of axonal and dendritic arborisations. These nascent contacts are believed to act like bolts during construction, yet a detailed understanding of their molecular composition and assembly is not known.

To explore this biology, we searched the fly for neurons that grow similarly to those in the fish and frog visual systems. Although fly embryos have been an excellent tool for studying connectivity, the small size and rapid development of their axonal arborisations makes it very challenging to image their growth live.

The axonal arborisations of the pleural muscle motoneurons (PM-Mns) are superficial and can be imaged *in vivo* throughout metamorphosis. Unlike the da sensory neuron input arborisations, which have been useful for studying neurite branch growth (Williams & Truman, 2004), PM-Mn axons form synapses; a trait in common with the majority of neuronal arborisations.

PM-Mn growth is very dynamic with a high turnover of exploratory filopodia, only a few of which ultimately become branches. Arborisations develop in close association with the pleural muscles and we find that axonal filopodia are stabilised upon contact with myoblasts or immature myotubes (see movie 2). A number of *in vitro* and *in vivo* studies have previously highlighted that filopodial stability is conferred by the contacts made between potential synaptic partners (Cooper & Smith, 1992; Ziv & Smith, 1996; Jontes *et al*., 2000).

To determine whether PM-Mn axonal arborisations could be growing using nascent synapses we imaged the active zone marker Bruchpilot (BRP) (Chen *et al*., 2014; Urwyler *et al.*, 2015) and vesicular machinery (VGlut and Syt1) and found similar localisations of synaptic proteins, as well as relationships with branch lifetimes as have been described in zebrafish and *Xenopus* (Alsina *et al*., 2001; Niell *et al*., 2004; Meyer & Smith, 2006; Ruthazer *et al*., 2006). BRP puncta appear to chart the progression of branch stabilisation events. To our surprise we also found Bruchpilot at similar localisations in the branches of developing eve+ interneurons in the CNS, indicating that this type of growth may be common within the fly nervous system.

### Growing arborisation do not require functional synapses

The exact contributions of synaptogenesis, neural activity and synaptic transmission to the formation of neural networks are unclear. Indeed, there is a rich history regarding the question of whether nervous systems develop in “*forward reference*” to, but without benefit from, functional activity (Harrison, 1904, Weiss,1941; see Haverkamp, 1986 for review).

Interestingly, we find that a great deal of PM-Mn outgrowth takes place prior to robust presynaptic calcium transients (action potentials, <42h APF). Furthermore, not until 60h APF could we evoke activity in muscles by stimulating the motoneurons. Finally, neither TTX or removing all vesicular neurotransmission, by making *VGlut* null clones, had an overt impact on mature arbor morphology. It may be that different systems are intrinsically different; Cline and colleagues showed a correlation between the stabilisation of growing retinal ganglion cell axon branches bearing synaptic puncta and visual activity, pointing to use-testing of nascent contacts by activity in a synaptotropic-like mode of growth (Ruthazer *et al*., 2006). In contrast however, a number of studies have shown that blocking activity has little impact on morphology, or plays only a role in the refinement of arbor growth (Haverkamp, 1986, Verhage *et al*., 2000; Varoquoaeux *et al*., 2002; Hua *et al*., 2005; Fredj *et al*., 2010).

The detailed imaging possible with our system reveals that BRP puncta very rarely enter filopodia and are never stabilised within them. We found the same to be true for the synaptic vesicle associated proteins Syt1 and VGlut. Since BRP is an important structural component of the cytomatrix at the active zone, these data seemed at odds with the notion that synapses drive branch stabilisation and led us to question if puncta previously reported at branch nodes in growing retinal ganglion cells really represent genuine synapses. Alongside this, we found it was not until late stages of development that the size and distribution of BRP puncta became more akin to synapses at the larval NMJ (Fouquet *et al*., 2009). In parallel, we saw conspicuous clusters of glutamate receptors only at late stages, long after arbor shape has been established. Taken together these data suggest that it is only at the later stages of arbor development that synaptic elements pair to form bona fide synapses. The accumulations of presynaptic proteins seen in early arborisations (i.e. 30-40h APF) may instead mark stable sites or ‘transport hubs’ that help store/sort synaptic proteins for a later role in synapse formation.

### Neuritic adhesion complexes (NACs) drive a tropic mode of arborisation growth

If synaptic transmission does not play a role in this type of dynamic arbor growth, what mechanisms are responsible? We find a role for a class of proteins (synaptic cell adhesion molecules), pre-synaptic Neurexins (Nrxs), and their postsynaptic binding partners, Neuroligins (Nlgs).

The *Nlg1* mutants and the clonal FlpStop data suggest that Nlg1 in the muscle targets is crucial for normal growth. Clonal analysis reveals that branches growing onto muscles with lower levels of *Nlg1* expression are phenotypically similar to those in *Nlg1* nulls. The growth of branches onto territories with more wild-type levels of *Nlg1* expression is reminiscent of the tropic aspect suggested in Vaughn’s original hypothesis.

Work in *Xenopus* has previously implicated Nrx-Nlg interactions in arbor growth and posited that synapse formation and subsequent levels of synaptic transmission translates directly into branch stability (Chen *et al*., 2010). Here we propose that these molecules provide adhesion during elaborative phases independent of synapse formation. We describe for the first time that Nlg1::GFP puncta emerge on muscles *in vivo* following contact with a presynaptic filopodia. Nlg1 localises onto the tips and shafts of filopodia and regulates stability. Filopodia that ‘capture’ such puncta are significantly longer lived than those that do not. Furthermore, time-lapse imaging revealed that Nlg1::GFP act like anchor points by marking the limits of filopodial retraction. This is very different to the growth of zebrafish and mouse neuromuscular junctions, where motoneurons grow between pre-patterned plaques of acetylcholine receptors, as if hopping between stepping stones ((Yang *et al*., 2001). Panzer *et al*., 2006; Jing *et al.*, 2009).

Presynaptically we found Nrx puncta on the tips of filopodia and along their lengths i.e. at sites that mirror those of Nlg1. The two other proteins we also find at such sites are the presynaptic membrane associated proteins Syd-1 and Liprin-α, both of which are known to complex with Nrx. Liprin-α and Syd-1 are found on the membrane and coalesce into dynamic puncta that localise to filopodia tips. Liprin-α puncta, appear to limit filopodial retraction, indicating that sites marked by this protein represent the presynaptic counterparts to the structural anchor points marked by Nlg1.

Previously, Syd-1 and Liprin-α have been found to play a key role in orchestrating synapse assembly by recruiting and retaining other synaptic proteins. Liprin-α, Syd-1 and Nrx puncta at these stages of exuberant growth are transient and do not appear to become ‘future synapses’. Their fluidity speaks to a role in branch morphogenesis rather than in a stepwise, clockwork assembly of synaptic machinery at a particular place.

Based on our observations, we propose that these dynamic complexes are composed of a subset of proteins that have been previously called ‘synaptic cell adhesion proteins’ or drivers of synapse formation. As a placeholder we suggest calling these puncta *neuritic adhesion complexes* or NACs to highlight their role in axon growth *prior to synapse formation.* That we see direct evidence for the formation of these NACs on filopodia themselves suggests that dynamic interactions between filopodia and the postsynaptic target are fundamental determinants of tree construction.

This early role for Nlg1/Nrx based NACs in arbor growth is supported by strong evidence that exploratory filopodia and branch tips are highly sensitive to changes in the early expression of postsynaptic *Nlg1*. Our gain-of-function myoblast/myotube stripes of clones seem to have a ‘hyper-stabilising’ effect on wild-type neurons and ultimately constrain growth. Moreover there is some indication of preferential growth into regions with elevated postsynaptic Nlg1 expression. Remarkably, we were very surprised to see wildtype PM-Mns grow past muscles, into novel territories and make contacts with class IV da sensory neurons ectopically expressing Nlg1.

### Physiological consequences of Nlg1 disruption

The precise role that Nrxs and Nlgs play in nervous system development has been broadly disputed. Previous work has shown the strong synaptogenic potential of these proteins by expressing them in HEK cells or on micropatterned substrates revealing them to be potent regulators of hemisynapse formation (Scheiffele *et al*., 2000; Dean *et al*., 2003; Graf *et al.*, 2004; Chih *et al*., 2005; Lee *et al*., 2010; Czöndör *et al*., 2013). Others show that mice lacking three Nlg homologues or all three α-Nrxs build brains with grossly normal cytoarchitectures and synapse densities (Missler *et al*., 2003; Varoqueaux *et al*., 2006) and posit that their role is solely to modulate synapse function. Roles for Nlg and Nrx in synapse formation have previously been described in the fly (Li *et al*., 2007; Banovic *et al*., 2010; Chen *et al*., 2012).

Teasing apart the function of such gene families in vertebrates can been difficult due to the multiple copies of these genes and their degeneracy with other proteins (Sugita *et al*., 2001; Ko *et al*., 2009; Siddiqui *et al*., 2010; Xu *et al*., 2010; Ko *et al*., 2011; Soler-Llavina *et al*., 2011). Here in this simpler system we show that Nlg1/Nrx based adhesion complexes play a key role in the early growth of axonal arborisations prior to the formation of functional synapses. In addition, we find that disrupting Nlg1/Nrx signalling has little effect on the timing of development and size of active zones, indicating that tree morphogenesis is uncoupled from the formation of synapses. One prediction from this is that disruption in functional connectivity would largely be due to disruptions in growth and morphology of the arborisation. An important observation that frames this is the outcome from ectopically expressing Nlg1 in class IV da sensory neurons. There we see the generation of a ‘synthetic connectivity’ on the normally ‘asynaptic’ da sensory neurites with the development of mature, differentiated presynaptic structures in later pupal stages i.e. synaptic vesicles and BRP labelled synapses.

In support of this, our calcium imaging reveals that the normal patterns of muscle activity are disrupted when Nlg1 is overexpressed, with changes in the frequency and spatial heterogeneity of calcium. Since elevated postsynaptic Nlg1 expression results in condensed arborisations, we propose that this is a result of irregular innervation. Additionally, the behavioural analysis shows that Nlg1 null adults had significant motor deficits. Taken together these data provide a link between an adhesion-based mechanism of arbor growth and synaptic connectivity which ultimately effects function.

## Conclusions

In this study we have demonstrated, for the first time outside of vertebrates, a dynamic ‘synaptotropic-like’ mode of arbor. In contrast to the mechanisms that have previously been proposed, we find no evidence that functional synapses drive this type of growth. Instead, our data point to dynamic NACs, composed of Neurexin, Neuroligin, Liprin-α and Syd-1, being important for stabilising filopodia during arborisation growth. Alongside this, our evidence showing the same subcellular localisations in *Drosophila* central neurons points toward this mode growth being a universal mechanism for building complex trees. It may be that Berry and colleagues’ emphasis on filopodia in their ‘synaptogenic filopodial theory’ was correct. It is an appealing idea that such mechanisms could construct axonal and dendritic arborisations of a broad spectrum of shapes and sizes by making only subtle changes to a ‘stick and grow’ algorithm. Exactly which proteins constitute these adhesion complexes and how they interact with the machinery for regulating cytoskeletal remodelling will be a focus of future investigation.

## Materials and Methods

### Fly stocks

Flies were reared on a standard yeast-cornmeal-molasses diet. For time-sensitive experiments flies were raised at 25°C, or at room temperature if precise staging was not required.

*OK371-GAL4; UAS-myr::GFP*

*OK371-GAL4; UAS-myr::GFP, UAS-BRP::RFP*

*OK371-GAL4; UAS-Lifeact::Ruby/UAS-CLIP170::GFP*

*VGlut(Trojan)-T2A-LexA, LexAop-myrGFP*

*VGlut(Trojan)-T2A-LexA, LexAop-myrGFP; Mef2-GAL4, UAS-mCD8::Cherry*

*VGlut(Trojan)-T2A-LexA, LexAop-myrGFP; Mef2-GAL4, UAS-myr::tdTomato*

*OK371-GAL4, UAS-mCD8::Cherry/BRP::GFP*

*OK371-GAL4, UAS-mCD8:Cherry/VGlut::GFP*

*OK371-GAL4, UAS-mCD8:Cherry/Syt1::GFP*

*RN2-Flp, Tub-FRT-CD2-FRTGal4, UAS-mCD8GFP/UAS-BRP::RFP*

*OK371-GAL4, UAS-mCD8:Cherry/GluRIIE::GFP OK371-GAL4, UAS-GCaMP6m*

*VGlut(Trojan)-T2A-LexA; Mef2-GAL4, UAS-GCaMP6M/LexAop-TRPA 1*

*VGlut^NMJX^-Gal4, UAS-mCD8::GFP, hsFlp^122^; FRT^40A^, Df(2L)VGlut^2^; UAS-myr::GFP*

*VGlut^NMJX^-Gal4, UAS-mCD8::GFP, hsFlp^122^; FRT^40A^, Df(2L)VGlut^2^/FRT^40A^; UAS-myr::GFP*

*Elav-GAL4, hsFlp^122^; FRT^40A^, Df(2L)vglut^2^/FRT^40A^; UAS-cytoplasmicGFP*

*VGlut(Trojan)-T2A-LexA, LexAop-myr::tdTomato; Mef2-GAL4, UAS-Nlg1::GFP*

*UAS-Nrx::GFP; OK371-GAL4, UAS-mCD8::Cherry*

*OK371-GAL4, UAS-mCD8::Cherry/UAS-Liprin-α::GFP*

*OK371-GAL4, UAS-Liprin-α::GFP/UAS-Syd 1::Straw*

*OK371-GAL4, UAS-Liprin-α::GFP; UAS-BRP::RFP*

*RN2-Flp, Tub-FRT-CD2-FRT-GAL4, UAS-mCD8::GFP/UAS-Liprin-α::GFP*

*VGlut(Trojan)-T2A-LexA, LexAop-myrGFP; Df(3R)BSC747/Nlg1^ex2.3^*

*VGlut(Trojan)-T2A-LexA, LexAop-myrGFP; Df(3R)BSC747/Nlg1^1960^*

*VGlut(Trojan)-T2A-LexA, LexAop-myrGFP; Df(3R)Exel6191/Nrx^241^*

*VGlut(Trojan)-T2A-LexA, LexAop-myrGFP; Mef2-GAL4/UAS-Nlg1^untagged^*

*hsFlp^122^; VGlut(Trojan)-T2A-LexA, LexAop-myrGFP, αTub84B-FRT.GAL80; Mef2-GAL4, UAS-myr::tdTomato/UAS-Nlg1^untagged^*

*PPK-GAL4, UAS-dsRed/VGlut-LexA, LexAop-myr::GFP*

*PPK-GAL4, UAS-dsRed/VGlut-LexA, LexAop-myr::GFP; UAS-Nlg1^untagged^*

*PPK-GAL4, UAS-mCD8::GFP; UAS-Nlg1^untagged^*

*VGlut(Trojan)-T2A-LexA, LexAop-myrGFP; Nlg1^FlpStop D^/Df(3R)BSC747*

*VGlut(Trojan)-T2A-LexA, LexAop-myrGFP; Nlg1^FlpStop ND^/Df(3R)BSC747*

*hsFlp^122^; VGlut(Trojan)-T2A-LexA, LexAop-myrGFP/Mef2-GAL4; Nlg1^FlpStop ND^/Df(3R)BSC747*

*hsFlp^122^; VGlut(Trojan)-T2A-LexA, LexAop-myrGFP/Mef2-GAL4; Nlg1^FlpStopD^/Df(3R)BSC747*

*Mef2-Gal4, UAS-GCaMP6m; UAS-Nlg1^untagged^*

*Mef2-Gal4, UAS-GCaMP6m*

### Imaging

Imaging was performed at room temperature using Zeiss LSM 510 or 800 series confocal microscopes with EC Plan-Neofluar 20x/0.50 or Plan-Apochromat 40x/1.3 objectives, except the laser ablation experiments, which were performed using a Nikon A1R with an Apochromat 40x/1.25 objective.

### Mounting and live imaging

For consistent developmental staging, pupae were collected at 0h APF (white prepupal stage) and incubated on moist tissue paper at 25°C in parafilm sealed petri dishes. In preparation for mounting, pupae were dried on tissue paper, immobilised on strips of double sided sticky tape and carefully removed from their puparial cases using forceps. Pupae were mounted using 22x22mm cover slips beneath a thin coating of halocarbon oil. For imaging sessions of greater than 1-hour pupae were mounted in humidity maintained chambers consisting of a platform of semi-permeable membrane suspended above a hole in a specialised steel slide and sealed with a watertight ring of petroleum jelly.

For experiments requiring temperature manipulation, a specialised temperature adjustable stage was built consisting of a glass slide fastened to a 5x5cm thermoelectric Peltier device (Maplin Electronics Ltd., UK) which was mounted on a regular microscope slide-holder. Before each experiment the temperature of the stage was calibrated using a thermocouple probe attached to a multi-meter device (Rapid Electronics Ltd., UK) and controlled from a 0-30v power supply unit.

### Mosaic analysis

The MARCM method was used to generate and label *VGlut* null motoneuron clones. A preliminary MARCM screen using OK371-GAL4 found that the pleural muscle motoneurons are born during the embryonic wave of neurogenesis. To generate clones during this wave of mitosis, breeding adults were allowed to lay on grape jelly plates with yeast for 2 hours at 25°C. Eggs were incubated for 3 hours at 25°C and heat shocked by incubation in a water bath at 37°C for 45 minutes, followed by resting at room temperature for 30 minutes, followed by a further 30 minutes at 37°C. Upon hatching, larvae were transferred to standard food and raised at 25°C.

To generate small numbers of GAL80 ‘flip-out’ or FlpStop Mef2-GAL4 positive muscle clones L3 wandering larvae were heat shocked for 20 minutes by incubation in a water bath at 37°C.

### Dissections and immunocytochemistry

Dissections were made in Sylgard silicone elastomer (Dow Corning, USA) lined petri dishes in 1x phosphate buffered saline (PBS) (pH 7.3, Thermo Fisher Scientific, USA).

For dissections of pupal abdominal body walls, pupae were removed from their puparial cases and pinned via the head using electrolytically sharpened 0.1mm tungsten pins. The posterior tip of each abdomen was removed (from approximately segment 6) and an incision made along the dorsal midline until reaching the thorax. Abdomens were separated from the thorax by cutting along the joint and laid flat using additional tungsten pins. Viscera were removed by a combination of gentle pipetting and forceps. Larval body wall dissections were performed as described previously (Broadie & Bate,1993)

Dissected samples were fixed in buffered 3.6% formaldehyde for 45 minutes at room temperature and washed three times in PBS containing 0.3 Triton-X100 (Sigma, USA) (PBST). Following fixation, samples were blocked in PBST containing 4% goat serum (Sigma, USA) for 45 minutes and incubated in solutions of primary antibodies made up with PBST overnight at 4°C. After rinsing 3 times with PBST over the course of a day, samples were incubated in solutions of secondary antibodies made up with PBST overnight at 4°C. After 3 rinses with PBST and a final rinse with PBS, samples were mounted on poly-L-lysine coated coverslips, dehydrated through a series of alcohol, cleared with xylene and mounted in DePeX (BDH Chemicals, UK).

Primary antibodies were used at the following concentrations: mouse anti-NC82 (DSHB, USA) 1:5, rabbit anti-VGlut-C term (Gift from Hermann Aberle (Mahr & Aberle, 2006)) 1:10,000, chicken anti-GFP (Invitrogen, USA) 1:1000. Secondary antibodies were used at the following concentrations: Cy3 goat anti-HRP (Jackson Immunoresearch, USA) 1:5000, AlexaFluor 488 goat anti-Chicken (Invitrogen, USA) 1:500, Cy5 donkey anti-mouse 1:500, Cy5 donkey antirabbit 1:500.

For dissections of CNSs, white pre-pupae were collected and incubated at 25°C. Pupae were dissected from their puparial cases as described above. CNSs were dissected out in 1x PBS in Sylgard lined Petri-dishes. These were then immediately fixed in 3.6% formaldehyde for 20 minutes. Fixative was washed off thoroughly using 1x PBS. CNSs were mounted on glass slides in Slow-Fade Antifade reagent (Thermo Fisher Scientific) and sealed beneath coverslips using nail-varnish.

### Microinjections

To prepare for injection, pupae were dried on tissue paper and secured to glass slides using double sided sticky tape. Needles, pulled from glass capillaries, were loaded with 0.5mM TTX made up in PBS or a control solution of PBS. Injections were made into the abdominal cavities through small access holes made in the puparial cases, using a Transjector-5246 micro-injector (Eppendorf, Germany). Following injection, animals were removed from the sticky tape with a damp paintbrush and incubated at 25°C.

### Climbing assay

Adult females were used two days post-eclosion. To assess climb speed, each fly was placed in a square plastic tube with 1 cm edges and transparent walls. The narrow calibre prevented flies from jumping or flying. Each fly was tapped to the bottom and its climb recorded with an S-PRI high-speed camera (AOS Technologies AG, Switzerland) at 700 fps, frame size of 900x700 pixels, under red light illumination. Automated body tracking was performed using FlyLimbTracker (Uhlmann *et al*., 2017) in Icy (de Chaumont *et al*., 2012) over a distance of 0.27 to 0.8 cm. Only recordings in which the climbing path had a linearity over 90%, as determined with the Motion Profiler processor in Icy, were considered for analysis. The average climb speed of each fly was determined by calculating the displacement of the centroid of the body between each frame over 3 repeats.

### Image Analysis

Raw image stacks were imported into the image processing software Fiji (http://imagei.net/Fiii) for enhancement of brightness and contrast. For clarity, the freehand select tool was used to remove obscuring objects created by degenerating larval tissues and macrophages on a slice by slice basis. Collapsed z-projections were imported into Photoshop (Adobe, USA) for figure assembly. Schematic cartoons were made in Photoshop or Illustrator (Adobe, USA). To generate kymographs the re-slice tool in FIJI was used to reconfigure the axis of image sequences.

### Statistical analysis

Statistical analyses were performed using the software package GraphPad Prism (GraphPad Software Inc., USA). Significances correspond to the following p values: <0.05 = *, <0.01 = **, <0.001 = ***, <0.0001 = **** For selecting between parametric and non-parametric tests, the residuals of each data set were tested for normality. Results are reported with standard deviations (SDs).

#### Quantification of synaptic puncta

Relationships between BRP::RFP, Nlg1::GFP and Liprin-α::GFP puncta dynamics and branch growth were calculated by manually counting and tracking puncta in FIJI. Puncta were identified by accumulations of fluorescent protein measuring 3 or more pixels in diameter at the native resolution. The diameters of BRP::RFP puncta were measured by intersection with the line tool in FIJI at their widest point and calculation of the full width at half maximum of the peak in fluorescence.

#### Morphometric analysis

Arbor reconstructions were generated using the semi-automated plugin for FIJI, Simple Neurite Tracer. Reconstructed skeletons were imported into a bespoke analysis software in which total arbor length, total branch number, Strahler order and total arbor were computed. Arbor area was calculated by giving branch coordinates a hypothetical ‘contact distance’ which represented its area of coverage. This value was set to 20*μ*m. In addition, this software could be used to add and delete vertices and nodes, allowing corrections to me made to tracing errors.

#### Straightness Index

To calculate branch bendiness, the ‘actual’ length of primary, secondary and tertiary axon branches was measured using the freehand line tool in FIJI as the distance between their nodes (or in the case of primary branches, between nodes and tips). These values were divided by the direct distances between these nodes, measured using the straight-line tool, giving an index of bendiness.

#### Active zone density

Active zone densities in axon terminals were calculated manually in FIJI using the multi-point tool. Bouton areas were calculated by tracing around boutons using the freehand selection tool.

## Acknowledgements

We would like to thank Hermann Aberle for providing the Nlg1 alleles and UAS reagents as well as the OK371-GAL4 enhancer-trap line and anti-VGlut antibodies, Stephan Sigrist for providing the Liprin-α and Syd1 reagents, Barret Pfeiffer for providing the UAS-GFP lines with translational enhancers, Aaron DiAntonio for sharing the *Df(2L)VGluŕ* allele, Frank Schnorrer for providing a copy of Mef2-GAL4, Vijay Raghavan for sending the FlyFos GluRIIE::GFP line and Tom Clandinin for kindly sharing his FlpStop construct and for advice on implementing it. In addition, we thank Jon Clarke for generously allowing us to use his confocal microscope. We also thank Matthias Landgraf for providing the VGlut-LexA, discussions and advice on experiments. Finally, we thank Jon Clarke, Matthias Landgraf, Martin Meyer and Laura Andreae for reading the manuscript. This work was funded by the BBSRC.

## Bibliography

Alsina, B., Vu, T. and Cohen-Cory, S. (2001). Visualizing synapse formation in arborizing optic axons in vivo: dynamics and modulation by BDNF. Nat. Neurosci. 4, 1093–101.

Andreae L. C. and Burrone J. (2015). Spontaneous Neurotransmitter Release Shapes Dendritic Arbors via Long-Range Activation of NMDA Receptors. Cell Rep. 10, 873–82.

Banovic, D., Khorramshahi, O., Owald, D., Wichmann, C., Riedt, T., Fouquet, W., Tian, R., Sigrist, S. J. and Aberle, H. (2010). Drosophila neuroligin 1 promotes growth and postsynaptic differentiation at glutamatergic neuromuscular junctions. Neuron 66, 724–38.

Bradley P. and Berry M. (1976). The growth of the dendritic trees of Purkinje cells in irradiated agranular cerebellar cortex. Brain Res. 35, 123–43.

Broadie K. and Bate M. (1993). Activity-dependent development of the neuromuscular synapse during Drosophila embryogenesis. Neuron 11, 607–19.

Chagnac-Amitai, Y., Luhmann, H. J. and Prince, D. a (1990). Burst Generating and Regular Spiking Layer-5 Pyramidal Neurons of Rat Neocortex Have Different Morphological Features. J. Comp. Neurol. 296, 598–613.

Chen S. X. and Haas K. (2011). Function directs form of neuronal architecture. Bioarchitecture 1, 2–4.

Chen, S. X., Tari, P. K., She, K. and Haas, K. (2010). Neurexin-neuroligin cell adhesion complexes contribute to synaptotropic dendritogenesis via growth stabilization mechanisms in vivo. Neuron 67, 967–83.

Chen, Y.-C., Lin, Y. Q., Banerjee, S., Venken, K., Li, J., Ismat, A., Chen, K., Duraine, L., Bellen, H. J. and Bhat, M. A. (2012). Drosophila neuroligin 2 is required presynaptically and postsynaptically for proper synaptic differentiation and synaptic transmission. J. Neurosci. 32, 16018–30.

Chen, Y., Akin, O., Nern, A., Tsui, C. Y. K., Pecot, M. Y. and Zipursky, S. L. (2014). Cell-type-specific labeling of synapses in vivo through synaptic tagging with recombination (STaR). Neuron 81, 280–93.

Chih, B., Engelman, H. and Scheiffele, P. (2005). Control of Excitatory and Inhibitory Synapse Formation by Neuroligins. Science (80-.). 307, 1324–8.

Choi, B. J., Imlach, W. L., Jiao, W., Wolfram, V., Wu, Y., Grbic, M., Cela, C., Baines, R. A., Nitabach, M. N. and McCabe, B. D. (2014). Miniature Neurotransmission Regulates Drosophila Synaptic Structural Maturation. Neuron 82, 618–34.

Cooper M. W. and Smith S. J. (1992). A real-time analysis of growth cone target-cell interactions during the formation of stable contacts between hippocampal-neurons in culture. J. Neurobiol. 23, 814–28.

Cuntz, H., Forstner, F., Borst, A. and Häusser, M. (2010). One rule to grow them all: A general theory of neuronal branching and its practical application. PLoS Comput. Biol. 6,.

Currie D. a and Bate M. (1991). The development of adult abdominal muscles in Drosophila: myoblasts express twist and are associated with nerves. Development 113, 91–102.

Czöndör, K., Garcia, M., Argento, A., Constals, A., Breillat, C., Tessier, B. and Thoumine, O. (2013). Micropatterned substrates coated with neuronal adhesion molecules for high-content study of synapse formation. Nat. Commun. 4,.

Daniels, R. W., Collins, C. a, Chen, K., Gelfand, M. V, Featherstone, D. E. and DiAntonio, A. (2006). A single vesicular glutamate transporter is sufficient to fill a synaptic vesicle. Neuron 49, 11–6.

de Chaumont, F., Dallongeville, S., Chenouard, N., Hervé, N., Pop, S., Provoost, T., Meas-Yedid, V., Pankajakshan, P., Lecomte, T., Le Montagner, Y., et al. (2012). Icy: an open bioimage informatics platform for extended reproducible research. Nat. Methods 9, 690–6.

Dean, C., Scholl, F. G., Choih, J., DeMaria, S., Berger, J., Isacoff, E. and Scheiffele, P. (2003). Neurexin mediates the assembly of presynaptic terminals. Nat. Neurosci. 6, 708–16.

Diao, F., Ironfield, H., Luan, H., Diao, F., Shropshire, W. C., Ewer, J., Marr, E., Potter, C. J., Landgraf, M. and White, B. H. (2015). Plug-and-play genetic access to drosophila cell types using exchangeable exon cassettes. Cell Rep. 10, 1410–21.

Fisher, Y. E., Yang, H. H., Isaacman-Beck, J., Xie, M., Gohl, D. M. and Clandinin, T. R. (2017). FlpStop, a tool for conditional gene control in Drosophila. Elife 6,.

Fouquet, W., Owald, D., Wichmann, C., Mertel, S., Depner, H., Dyba, M., Hallermann, S., Kittel, R. J., Eimer, S. and Sigrist, S. J. (2009). Maturation of active zone assembly by Drosophila Bruchpilot. J. Cell Biol. 186, 129–45.

Fredj, N. B., Hammond, S., Otsuna, H., Chien, C.-B., Burrone, J. and Meyer, M. P. (2010). Synaptic activity and activity-dependent competition regulates axon arbor maturation, growth arrest, and territory in the retinotectal projection. J. Neurosci. 30, 10939–51.

Graf, E. R., Zhang, X., Jin, S. X., Linhoff, M. W. and Craig, A. M. (2004). Neurexins induce differentiation of GABA and glutamate postsynaptic specializations via neuroligins. Cell 119, 1013–26.

Grueber, W. B., Ye, B., Moore, A. W., Jan, L. Y. and Jan, Y. N. (2003). Dendrites of Distinct Classes of Drosophila Sensory Neurons Show Different Capacities for Homotypic Repulsion. Curr. Biol. 13, 618–26.

Haas, K., Li, J. and Cline, H. T. (2006). AMPA receptors regulate experience-dependent dendritic arbor growth in vivo. Proc. Natl. Acad. Sci. U. S. A. 103, 12127–31.

Hamada, F. N., Rosenzweig, M., Kang, K., Pulver, S. R., Ghezzi, A., Jegla, T. J. and Garrity, P. A. (2008). An internal thermal sensor controlling temperature preference in Drosophila. Nature 454, 217–22.

Harrison R. G. (1902). An experimental study of the relationship of the nervous system to the developing musculature in the embryo of the frog. Am. J. Anat. 11, 197–220.

Hassan B. A. and Hiesinger P. R. (2015). Beyond Molecular Codes: Simple Rules to Wire Complex Brains. Cell 163, 285–91.

Haverkamp J. (1986). Anatomical and Physiological Development of the Xenopus Embryonic Motor System in the Absence of Neural Activity. J. Neurosci. 6, 1338–48.

Hossain, S., Sesath Hewapathirane, D. and Haas, K. (2012). Dynamic morphometrics reveals contributions of dendritic growth cones and filopodia to dendritogenesis in the intact and awake embryonic brain. Dev. Neurobiol. 72, 615–27.

Hua, J. Y., Smear, M. C., Baier, H. and Smith, S. J. (2005). Regulation of axon growth in vivo by activity-based competition. Nature 434, 1022–26.

Jing, L., Lefebvre, J. L., Gordon, L. R. and Granato, M. (2010). Wnt signals organize synaptic prepattern and axon guidance through the zebrafish unplugged/MuSK receptor. Neuron 61, 721–733.

Jontes, J. D., Buchanan, J. and Smith, S. J. (2000). Growth cone and dendrite dynamics in zebrafish embryos: early events in synaptogenesis imaged in vivo. Nat. Neurosci. 3, 231–7.

Kaethner R. J. and Stuermer C. a (1992). Dynamics of terminal arbor formation and target approach of retinotectal axons in living zebrafish embryos: a time-lapse study of single axons. J. Neurosci. 12, 3257–71.

Ko, J., Fuccillo, M. V., Malenka, R. C. and Südhof, T. C. (2009). LRRTM2 Functions as a Neurexin Ligand in Promoting Excitatory Synapse Formation. Neuron 64, 791–8.

Ko, J., Soler-Llavina, G. J., Fuccillo, M. V., Malenka, R. C. and Südhof, T. C. (2011). Neuroligins/LRRTMs prevent activity - and Ca 2+/calmodulin-dependent synapse elimination in cultured neurons. J. Cell Biol. 194, 323–34.

Lee, H., Dean, C. and Isacoff, E. (2010). Alternative Splicing of Neuroligin Regulates the Rate of Presynaptic Differentiation. J. Neurosci. 30, 11435–46.

Li, J., Ashley, J., Budnik, V. and Bhat, M. A. (2007). Crucial Role of Drosophila Neurexin in Proper Active Zone Apposition to Postsynaptic Densities, Synaptic Growth, and Synaptic Transmission. Neuron 55, 741–55.

Lye, C. M., Naylor, H. W. and Sanson, B. (2014). Subcellular localisations of the CPTI collection of YFP-tagged proteins in Drosophila embryos. Development 141, 40064017.

Mahr A. and Aberle H. (2006). The expression pattern of the Drosophila vesicular glutamate transporter: A marker protein for motoneurons and glutamatergic centers in the brain. Gene Expr. Patterns 6, 299–309.

Mattila P. K. and Lappalainen P. (2008). Filopodia: molecular architecture and cellular functions. Nat. Rev. Mol. Cell Biol. 9, 446–54.

Meyer M. P. and Smith S. J. (2006). Evidence from in vivo imaging that synaptogenesis guides the growth and branching of axonal arbors by two distinct mechanisms. J. Neurosci. 26, 3604–14.

Missler, M., Zhang, W., Rohlmann, A., Kattenstroth, G., Hammer, R. E., Gottmann, K. and Südhof, T. C. (2003). Alpha-neurexins couple Ca2+ channels to synaptic vesicle exocytosis. Nature 423, 939–48.

Niell, C. M., Meyer, M. P. and Smith, S. J. (2004). In vivo imaging of synapse formation on a growing dendritic arbor. Nat. Neurosci. 7, 254–60.

Owald, D., Fouquet, W., Schmidt, M., Wichmann, C., Mertel, S., Depner, H., Christiansen, F., Zube, C., Quentin, C., Körner, J., et al. (2010). A Syd-1 homologue regulates pre - and postsynaptic maturation in Drosophila. J. Cell Biol. 188, 565–79.

Owald, D., Khorramshahi, O., Gupta, V. K., Banovic, D., Depner, H., Fouquet, W., Wichmann, C., Mertel, S., Eimer, S., Reynolds, E., et al. (2012). Cooperation of Syd-1 with Neurexin synchronizes pre-with postsynaptic assembly. Nat. Neurosci. 15, 1219–26.

Panzer, J. A., Song, Y. and Balice-gordon, R. J. (2006). In Vivo Imaging of Preferential Motor Axon Outgrowth to and Synaptogenesis at Prepatterned Acetylcholine Receptor Clusters in Embryonic Zebrafish Skeletal Muscle. J. Neurosci. 26, 934–47.

Rajan, I., Witte, S. and Cline, H. T. (1999). NMDA receptor activity stabilizes presynaptic retinotectal axons and postsynaptic optic tectal cell dendrites in vivo. J. Neurobiol. 38, 357–68.

Ranganayakulu, G., Schulz, R. A. and Olson, E. N. (1996). Wingless Signaling Induces nautilus Expression in the Ventral Mesoderm of the Drosophila Embryo. Dev. Biol. 176, 143–8.

Riedl, J., Crevenna, A. H., Kessenbrock, K., Yu, J. H., Neukirchen, D., Bista, M., Bradke, F., Jenne, D., Holak, T. A., Werb, Z., et al. (2010). Lifeact: a versatile marker to visualize F-actin. Nat. Methods 5, 605–7.

Roberts, A., Conte, D., Hull, M., Merrison-Hort, R., al Azad, A. K., Buhl, E., Borisyuk, R. and Soffe, S. R. (2014). Can Simple Rules Control Development of a Pioneer Vertebrate Neuronal Network Generating Behavior? J. Neurosci. 34, 608–21.

Roy, B., Singh, A. P., Shetty, C., Chaudhary, V., North, A., Landgraf, M., VijayRaghavan, K. and Rodrigues, V. (2007). Metamorphosis of an identified serotonergic neuron in the Drosophila olfactory system. Neural Dev. 2,.

Ruthazer, E. S., Akerman, C. J. and Cline, H. T. (2003). Control of Axon Branch Dynamics by Correlated Activity in Vivo. Science (80-.). 301, 66–70.

Ruthazer, E. S., Li, J. and Cline, H. T. (2006). Stabilization of axon branch dynamics by synaptic maturation. J. Neurosci. 26, 3594–603.

Sarov, M., Barz, C., Jambor, H., Hein, M. Y., Schmied, C., Suchold, D., Stender, B., Janosch, S., Vinay Vikas, K. J., Krishnan, R. T., et al. (2016). A genome-wide resource for the analysis of protein localisation in Drosophila. Elife 5, 1–38.

Scheiffele, P., Fan, J., Choih, J., Fetter, R. and Serafini, T. (2000). Neuroligin expressed in nonneuronal cells triggers presynaptic development in contacting axons. Cell 101, 657–69.

Schnütgen, F., Doerflinger, N., Calléja, C., Wendling, O., Chambon, P. and Ghyselinck, N. B. (2003). A directional strategy for monitoring Cre-mediated recombination at the cellular level in the mouse. Nat. Biotechnol. 21, 562–5.

Siddiqui, T. J., Pancaroglu, R., Kang, Y., Rooyakkers, A. and Marie, A. (2010). LRRTMs and Neuroligins Bind Neurexins with a Differential Code to Cooperate in Glutamate Synapse Development. Brain Res. 30, 7495–506.

Sin, W. C., Haas, K., Ruthazer, E. S. and Cline, H. T. (2002). Dendrite growth increased by visual activity requires NMDA receptor and Rho GTPases. Nature 419, 475–80.

Soler-Llavina, G. J., Fuccillo, M. V, Ko, J. and Malenka, R. C. (2011). The neurexin ligands, neuroligins and leucine-rich repeat transmembrane proteins, perform convergent and divergent synaptic functions in vivo. Proc. Natl. Acad. Sci. U. S. A. 108, 16502–9.

Stramer, B., Moreira, S., Millard, T., Evans, I., Huang, C. Y., Sabet, O., Milner, M., Dunn, G., Martin, P. and Wood, W. (2010). Clasp-mediated microtubule bundling regulates persistent motility and contact repulsion in Drosophila macrophages in vivo. J. Cell Biol. 189, 681–9.

Sugita, S., Saito, F., Tang, J., Satz, J., Campbell, K. and Südhof, T. C. (2001). A stoichiometric complex of neurexins and dystroglycan in brain. J. Cell Biol. 154, 435–45.

Teeter C. M. and Stevens C. F. (2011). A general principle of neural arbor branch density. Curr. Biol. 21, 2105–8.

Uhlmann, V., Ramdya, P., Delgado-Gonzalo, R., Benton, R. and Unser, M. (2017). FlyLimbTracker: An active contour based approach for leg segment tracking in unmarked, freely behaving Drosophila. PLoS One 12,.

Urwyler, O., Izadifar, A., Dascenco, D., Petrovic, M., He, H., Ayaz, D., Kremer, A., Lippens, S., Baatsen, P., Guérin, C. J., et al. (2015). Investigating CNS synaptogenesis at single-synapse resolution by combining reverse genetics with correlative light and electron microscopy. Development 142, 394–405.

Varoqueaux, F., Sigler, A., Rhee, J.-S., Brose, N., Enk, C., Reim, K. and Rosenmund, C. (2002). Total arrest of spontaneous and evoked synaptic transmission but normal synaptogenesis in the absence of Munc13-mediated vesicle priming. Proc. Natl. Acad. Sci. U. S. A. 99, 9037–42.

Varoqueaux, F., Aramuni, G., Rawson, R. L., Mohrmann, R., Missler, M., Gottmann, K., Zhang, W., Südhof, T. C. and Brose, N. (2006). Neuroligins Determine Synapse Maturation and Function. Neuron 51, 741–54.

Vaughn J. E. (1989). Fine structure of synaptogenesis in the vertebrate central nervous system. Synapse 3, 255–85.

Vaughn, J. E., Henrikson, C. K. and Grieshaber, J. A. (1974). A quantitative study of synapses on motor neuron dendritic growth cones in developing mouse spinal cord. J. Cell Biol. 60, 664–72.

Vaughn, J. E., Barber, R. P. and Sims, T. J. (1988). Dendritic development and preferential growth into synaptogenic fields: a quantitative study of Golgi-impregnated spinal motor neurons. Synapse 2, 69–78.

Venken, K. J. T., Schulze, K. L., Haelterman, N. A., Pan, H., He, Y., Evans-Holm, M., Carlson, J. W., Levis, R. W., Spradling, A. C., Hoskins, R. A., et al. (2011). MiMIC: a highly versatile transposon insertion resource for engineering Drosophila melanogaster genes. Nat. Methods 8, 737–743.

Verhage, M., Maia, A. S., Plomp, J. J., Brussaard, A. B., Heeroma, J. H., Vermeer, H., Toonen, R. F., Hammer, R. E., van den Berg, T. K., Missler, M., et al. (2000). Synaptic assembly of the brain in the absence of neurotransmitter secretion. Science (80-.). 287, 864–9.

Walter, J., Henke-fahle, S. and Bonhoeffer, F. (1987). Avoidance of posterior tectal membranes by temporal retinal axons. Development 913, 909–13.

Weiss P. (1941). Self-differentiation of the basic patterns of coordination. Comp. Psychol. Monogr. 17, 1–96.

Wu, G. Y., Zou, D. J., Rajan, I. and Cline, H. (1999). Dendritic dynamics in vivo change during neuronal maturation. J. Neurosci. 19, 4472–83.

Xu, J., Xiao, N. and Xia, J. (2010). Thrombospondin 1 accelerates synaptogenesis in hippocampal neurons through neuroligin 1. Nat. Neurosci. 13, 22–24.

Yang, X., Arber, S., William, C., Li, L., Tanabe, Y., Jessell, T. M., Birchmeier, C. and Burden, S. J. (2001). Patterning of muscle acetylcholine receptor gene expression in the absence of motor innervation. Neuron 30, 399–410.

Ziv N. E. and Smith S. J. (1996). Evidence for a role of dendritic filopodia in synaptogenesis and spine formation. Neuron 17, 91–102.

